# A Near Infrared Fluorescent Nanosensor for Spatial and Dynamic Measurements of Auxin, Indole-3-Acetic Acid, *in Planta*

**DOI:** 10.1101/2024.05.23.595494

**Authors:** Duc Thinh Khong, Kien Van Vu, Benny Jian Rong Sng, Ian Kin Yuen Choi, Thomas K. Porter, Jianqiao Cui, Xun Gong, Song Wang, Nguyen Hoai Nguyen, Mervin Ang, Minkyung Park, Tedrick Thomas Salim Lew, Suh In Loh, Riza Ahsim, Hui Jun Chin, Gajendra Pratap Singh, Mary B. Chan-Park, Nam-Hai Chua, Michael S. Strano, In-Cheol Jang

## Abstract

Auxins, particularly indole-3-acetic acid (IAA), is a phytohormone critical for plant growth, development, and response to environmental stimuli. Despite its importance, there is a lack of species-independent sensors that allow direct and reversible detection of IAA. Herein, we introduce a novel near infrared fluorescent nanosensor for spatial and temporal measurement of IAA *in planta* using Corona Phase Molecular Recognition. The IAA nanosensor shows high specificity to IAA *in vitro* and was validated to localize and function in plant cells. The sensor works across different plant species without optimization and allows visualization of dynamic changes to IAA distribution and movement in leaf tissues. The results highlighted the utility of IAA nanosensor for understanding IAA dynamics *in planta*.

## Introduction

As climate change stresses global agricultural production ^1^, there is a pressing need for deeper understanding of the molecular processes inside plants that regulate stress responses, which ultimately affect crop yield and quality ^2^. Central to these processes is indole-3-acetic acid (IAA), a naturally occurring auxin that regulates plant physiological processes^3^ [NO_PRINTED_FORM] and environmental responses such as gravitropism, phototropism, hydrotropism, and shade avoidance ^4^. At the cellular level, the temporal and gradient distribution of IAA across plant cells affects cellular growth and behavior such as cell elongation, division, expansion, and differentiation. Auxin activity is highly regulated by its biosynthesis and transport ^5,6^, and its signaling influences the plant response to environmental stressors ^4,7,8^. However, it has been a long-standing challenge to develop effective methods for real-time and spatial-temporal visualization of IAA *in planta* to elucidate persistent questions about IAA activity.

Several techniques that evaluate the signature or trace of IAA distribution in plants have been reported in the literature ^9^. Conventional methods for sensing IAA were either indirect, basing on the expression of auxin-responsive genes such as *AUXIN/INDOLE-3-ACETIC ACID* (*Aux/IAA*), *GRETCHEN HAGEN 3* (*GH3*), and *SMALL AUXIN-UP RNA* (*SAUR*) gene families ^10^, or direct quantification of IAA with mass spectrometry ^11^. These methods involve homogenization of plant tissues and extraction of RNA transcripts or metabolites, thus lacking spatial and temporal resolution. However, to study the many downstream developmental processes including embryogenesis, lateral root formation, development of initiation sites in the shoot apical meristem and various tropisms ^12^, it is crucial to understand the spatial-temporal distribution of auxin along with its gradient and local maxima ^3^. The transgenic reporter systems are commonly utilized for visualizing auxin distribution *in planta*, achieved through the expression of visual markers such as β-glucuronidase (GUS) or green fluorescent protein (GFP) under the control of synthetic auxin-responsive DR5 promoter (*DR5::GUS)* ^13^ or *DR5::GFP* ^14^. Auxin distribution can also be inferred by monitoring the auxin-responsive protein degradation of AUX/IAA domain II, which is fused with VENUS yellow fluorescent protein (*35S::DII-VENUS*) ^15^. Recently, a genetically encoded fluorescent biosensor based on Fluorescence Resonance Energy Transfer (FRET) has been developed for direct IAA visualization. Engineered from *Escherichia coli* tryptophan repressor protein modified with an IAA-specific binding pocket, this sensing system shows direct spatiotemporal observation of auxin levels in the root apex and its transient changes during exogenous auxin uptake ^16^. However, like other genetically encoded biosensors, their development requires generation of transgenic plants, which are time-consuming and restricting its use to specific plant models with established methods for genetic engineering ^17^. Furthermore, the optical range of these fluorescent biosensors are typically in the visible spectrum region, which gives sub-optimal *in vivo* imaging because of weaker tissue penetration, high scattering, and confounding signals from cellular autofluorescence ^18^. Therefore, while live imaging of *in planta* auxin distribution was mostly carried out in root and young tissues such as shoot apical meristem or leaf initiation ^19–23^, visualization of IAA in the matured leaves is limited due to strong interference from chlorophyll autofluorescence ^24,25^.

To address these limitations, we employed plant nanobionics technology ^26^ to develop a new sensing platform using nanoparticles for high-resolution analysis of spatiotemporal distribution of IAA in living plants at stand-off distance The IAA nanosensor was developed using a synthetic construct that recognizes IAA and attaches to carbon nanotubes for near-infrared (nIR) fluorescence detection. Such nanosensor probes have advantagous optical properties for in-vivo chemical sensing. They are non-photobleaching, and possess narrow emission throughout the nIR region, avoiding biological autofluorescence^27^. Unlike transgenic methods, our synthetic nanosensor can be readily infused into living plants, providing a robust and flexible approach for real-time observation of IAA distribution ^28,29^. Our nanosensor uses a similar sensing platform as Corona Phase Molecular Recognition (CoPhMoRe) ^30^ in its development, yet it is distinct from other constructs for its target analyte ^28^. In this work, we demonstrate that the CoPhMoRe-based IAA nanosensor can be effectively introduced into plant cells and enables real-time imaging of IAA concentration and distribution in the leaf tissues of *XVE::iaaM* Arabidopsis plants, effectively showing the induced biosynthesis and local transport of IAA *in planta*. Our results illustrate the potential of this sensor platform to further uncover the spatial-temporal importance of IAA in regulating plant growth and development, as well as to serve as a diagnostic tool for early detection of IAA-related plant stresses that hinder global agriculture ^31^.

## Results

### Design of IAA nanosensor by taking cues from the IAA interaction with TRANSPORT INHIBITOR RESPONSE1 (TIR1)

The IAA nanosensor was developed based on the CoPhMoRe platform that we have described previously ^30^. This approach involves the physical adsorption of a synthetic polymer on single wall carbon nanotube (SWNT) surface, creating an aqueous corona phase that allows for selective docking of the IAA analyte. The synthetic corona phase for IAA sensing was designed, screened, and optimized from a compositionally varied library of synthetic polymers. By employing carbon nanotube as the signal transducer, the sensor offers highly photostable bandgap nIR fluorescence with wavelengths that are often non-interfering with photo-absorption or autofluorescence windows of living tissue ^32^. Our design for IAA sensing was inspired from the known interactions between IAA and the TRANSPORT INHIBITOR RESPONSE1 (TIR1) ^33^. TIR1 is an F-box protein subunit of the ubiquitin ligase complex (SCF^TIR1^) that is primarily involved in auxin-mediated signaling pathways ^34,35^. Crystallographic study on the TIR1-IAA complex reveals several key binding modes, including: (i) the salt-bridge interaction between IAA carboxyl group and a guanidium residue (Arg 403) of TIR1, coupled with a hydrogen bonding of IAA carboxyl group to another hydroxyl residue of TIR1 serine building block (Ser 438), (ii) the hydrophobic and van der Waals interaction between IAA indolic group and phenyl residues (Phe 79 and Phe 82) on TIR1, and (iii) the hydrogen bonding between indolic nitrogen of IAA and a carbonyl on the side of TIR1 binding pocket ^33^ (Fig. 1a). To capture the interactions between hydrophilic and hydrophobic portions of IAA, we proposed the use of synthetic amphiphilic polyamic sodium salts with tunable functional groups as the backbone of corona phase construct.

**Fig. 1.**
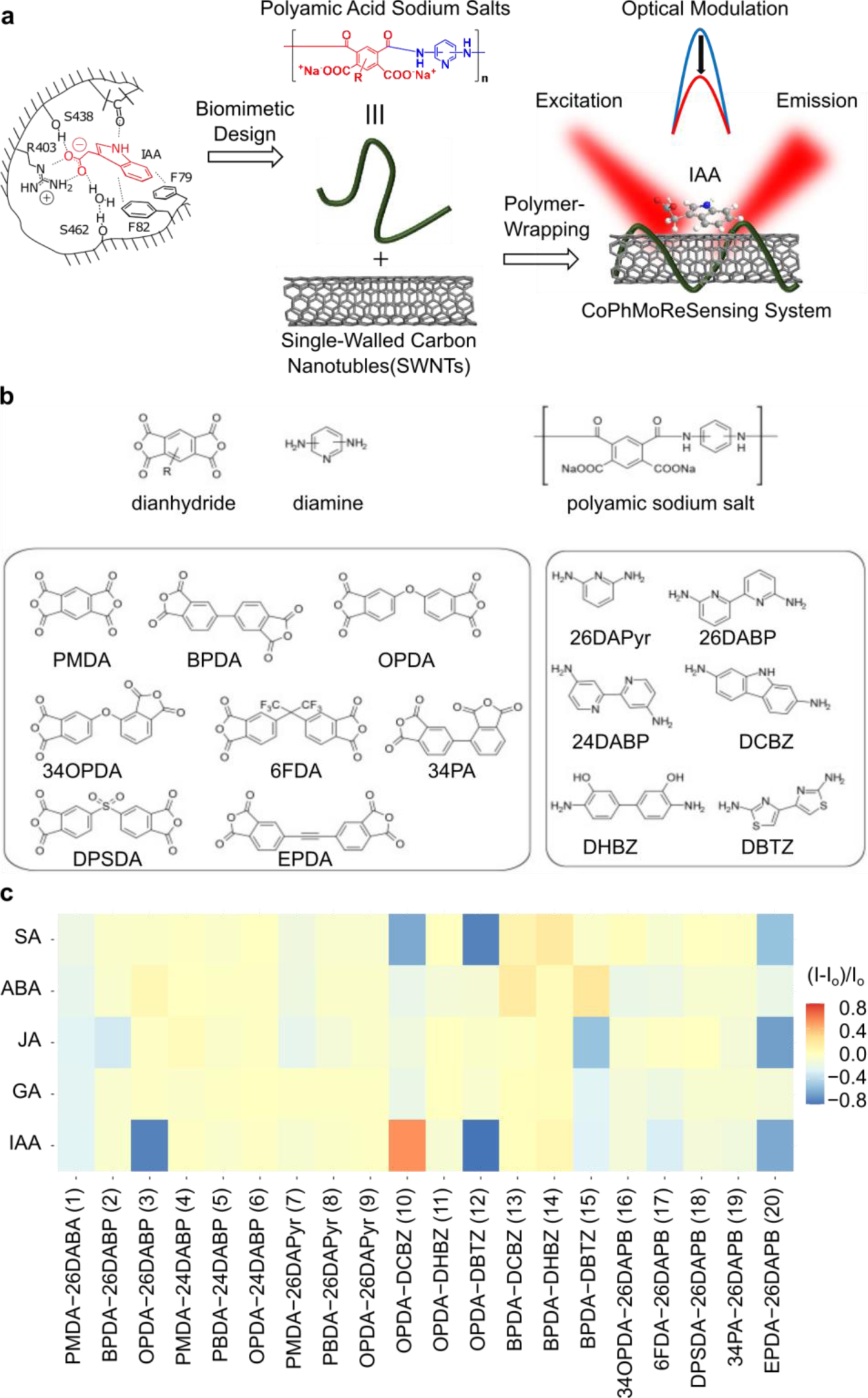
Design, Development, and Screening of IAA Nanosensor. **(a)** Schematic diagram illustrating the semi-rational design of the CoPhMoRe IAA sensor based on the in vivo interaction between IAA and the TIR1 binding pocket, resulting in fluorescent modulation upon IAA binding. **(b)** Synthetic scheme for polyamic acid sodium salts. A library of 20 polyamic acid sodium salts was synthesized using combinations of different aromatic dianhydrides and diamines (see Supplementary Fig. 1 and 2 for complete molecular structures of monomers and polymer library). Bottom left rectangular box: List of dianhydride monomers used in synthesis. Bottom right rectangular box: List of diamine monomers used in synthesis. **(c)** Heatmap summarizing the nIR fluorescent responses of CoPhMoRe sensors (1)−(20) (x-axis) to plant analytes (y-axis). Fluorescent intensity was measured before and after the addition of plant analyte (100 µM) using a home-built stand-off nIR camera setup with an excitation wavelength of 785 nm and laser power of 30 mW. Normalized fluorescent intensity changes (I-I_o_)/I_o_ are shown in a color-coded heatmap where negative values indicate quenching response and positive values indicate turn-on response. Polymer structures of CoPhMoRes are provided in supplementary information. Data in each square represents the mean normalized intensity change obtained from three independent measurements (*n*=3).

### Synthesis of CoPhMoRes library for IAA nanosensor screening

In general, polyamic acid sodium salts are neutralized water-soluble derivations of polyamic acids, which are a type of co-polymer system formed by condensation reaction between an aromatic dianhydride and aromatic diamine. The backbone of these polymers is rich with phenyl moieties which is known for π-π stacking on the nanotube surface ^36^, assisting the formation of stable aqueous corona phase for potential IAA docking. It also has a high density of amide bonds, carboxylate/carboxylic groups and pyridine nitrogen-based functional groups which can be modified for preferential hydrogen bonding and ionic interactions with the corresponding functional group in IAA (Fig. 1a-b). Based on the corresponding list of synthesized polyamic acids sodium salts (Fig. S1, S2), a library of 20 CoPhMoRe-based sensors for IAA sensing was generated, through non-covalent conjugation with SWNTs using the tip-sonication method ^30^. Characterization of these CoPhMoRes with UV-Vis-nIR spectroscopy showed well-defined absorption chirality peaks in the S_11_ and S_22_ regions, indicating that these polymers wrap non-selectively to all SWNT chiralities with suspension yields up to 50−200 mg L^−1^ (Fig. S3).

### Screening of nanosensor candidates against IAA and other phytohormones

Screening was carried out using photoluminescence spectroscopy to investigate the nIR optical modulation of each CoPhMoRe sensor against IAA and other phytohormones. The evaluation of optical modulation is based on the normalized fluorescence intensity changes, (I-I_o_)/I_o_, before (I_o_) and after (I) analyte addition. A false color-coded heatmap in Fig. 1c summarizes the optical response of the 20 CoPhMoRe sensors against IAA and other phytohormones (e.g., GA_3_, ABA, JA, SA). We observed that CoPhMoRe sensor #3, made from two monomers (OPDA and 26DAPB), selectively and strongly responded to 100 μM IAA with 75% fluorescence quenching while being insensitive to other analytes (Fig. 1c). Other CoPhMoRe sensors were not selected because of weak response or low specificity to IAA (Fig. 1c). In addition, both IAA in deprotonated (neutralized by NaOH) and protonated (diluted in DMSO) forms responded well to CoPhMoRe #3 with 75% and 52% quenching, respectively (Fig. 2a). The weaker quenching response in protonated form may be due to competitive quenching by DMSO (Fig. 2a).

**Fig. 2.**
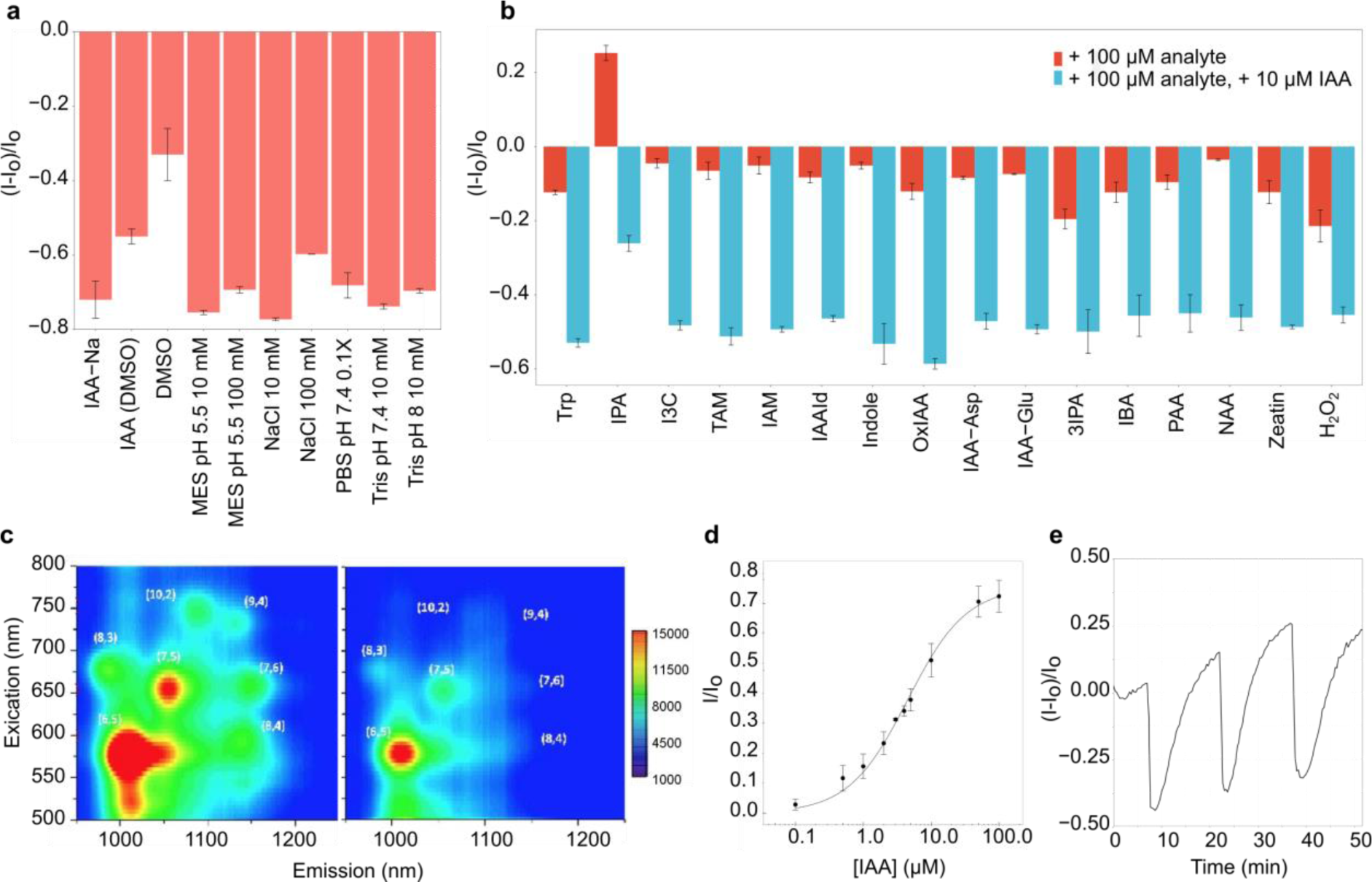
*In vitro* investigation on the IAA nanosensor. **(a)** Optical response of CoPhMoRe #3 to IAA in different ionic states (first two columns) and aqueous buffer conditions. **(b)** Interference of IAA sensor by compounds with similar structure or physiological function in 10 mM MES pH 5.5 buffer. Red columns: Sensor response (I-I_o_)/I_o_ after addition of IAA analogs and other related metabolites. Cyan columns: Sensor response (I-I_o_)/I_o_ after addition of 100 µM IAA to solutions already containing interfering compounds. Sensor response represents the final response in the presence of both IAA and interfering molecules. **(c)** Excitation-emission profile showing in vitro response of IAA sensor to 100 μM IAA. The sensor exhibits a reduction in fluorescent intensity for all observed chiralities (designated as pair number (n, m) on the heatmap). Estimation of dissociation constants for each nanotube chirality was carried out using excitation wavelengths of 750 nm for (10,2) and (9,4), 650 nm for (7,5) and (7,6), and 600 nm for (6,5) and (8,4). **(d)** Calibration curve of IAA sensor response I/Io to IAA. The dashed fitting curve was plotted based on the Langmuir adsorption model. **(e)** Reversible IAA response (I-I_o_)/I_o_ to IAA sensor demonstrated by in situ IAA photodegradation. For **a**, **b**, and **d**, data is present as mean ± s.d. from three independent experiments (*n*=3). For **c** and **e,** three independent experiments were repeated with similar results (*n*=3). Trp: L-tryptophan, IPA: indole-3-pyruvic acid, I3C: indole-3-carbinol, IAM: indole-3-acetamide, IAAId: indole-3-acetaldehyde, OxIAA: 2-oxindole-3-acetic acid, IAA-Asp: indole-3-acetyl-aspartate, IAA-Glu: indole-3-acetyl-glutamate, 3IPA: indole-3-propionic acid, IBA: indole-3-butyric acid, PAA: phenylacetic acid, NAA: 1-naphthaleneacetic acid.

### In vitro investigation of IAA nanosensor

The selected IAA nanosensor candidate (CoPhMoRe #3) was evaluated for its performance in various aqueous environments and found to respond similarly in physiological pH conditions (5.5-8) and ionic strength (10-100 mM NaCl) with an average of 71%±5% fluorescence quenching to 100 µM IAA (Fig. 2a). For further investigation on its selectivity and specificity, the sensor responses were measured against a wide range of potential interfering analytes at 100 µM concentration, including auxin homologs, auxin-related metabolites and other relevant plant signaling molecules (Fig. 2b, red columns). The results showed that IAA sensor was relatively inert in the presence of those analytes, except for IPA – a biosynthetic precursor to IAA, which showed a 33% increase in fluorescence intensity (Fig. 2b). In general, the interaction between IAA and the sensor was still preferential compared to other IAA-related analytes, as the sensor displayed approximately 50% fluorescence quenching despite only 10 µM IAA was added to the sensor solutions containing 100 µM of potential interfering analytes (Fig. 2b, cyan columns). As for IPA, further addition of 10 µM IAA resulted in 25% quenching of the sensor fluorescence (Fig. 2b, cyan columns).

To investigate the binding dynamics between IPA and IAA, a fluorescence kinetic study was conducted. The results, shown in Fig. S4, reveal that upon the addition of IPA, the fluorescence intensity quickly increased, but eventually returned to a quenched state in the presence of both IPA and IAA at 10:1 molar ratio. This indicates that while IPA binds to the sensor rapidly, IAA binding is thermodynamically favored, resulting in the eventual return of the fluorescence to a quenched state at equilibrium.

### *In vitro* calibration and reversibility of IAA nanosensor

The excitation-emission heatmap of IAA nanosensor in response to 100 µM IAA demonstrated that all observed SWNT chiralities underwent fluorescence quenching upon IAA addition (Fig. 2c). This suggests that IAA does not exhibit any preferential interaction with SWNT chirality. A calibration curve for the IAA sensor was generated *in vitro*, with IAA concentrations ranging from 0.01 to 100 µM, and showed that the sensor could detect IAA at concentrations as low as 0.1 µM (Fig. 2d). This is within the range of *in planta* detection, as the average endogenous IAA concentration in Arabidopsis has been reported to be approximately 0.1-3 µM ^37^. The dissociation constant coefficient K_d_, which measures the binding affinity of IAA to the sensor, was estimated using Langmuir adsorption isotherm model:

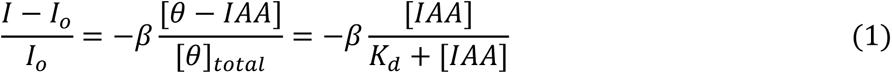

Where *β* is proportional factor showing the maximum optical modulation, [*θ* − *IAA*] and [*θ*_*total*_] are the concentration of occupied and total binding sites respectively. By fitting the experimental data into equation (1), the general K_d_ was estimated to be 4.5±0.21 μM. Estimation of specific K_d_ and *β* values for each SWNT chirality showed a slight dependence on chirality type to IAA affinity (Fig. S5). SWNT chirality (10,2) seems to have least affinity to IAA with dissociation constant *K_d_* = 4.33 ± 0.51 μM while the other three, (7,6), (7,5) and (8,4), have higher affinities with *K_d_* ≅ 1.2–1.5 μM. The data suggests that using chirality-separated SWNTs such as (7,6) could potentially yield more refined fluorescent modulation (by exclusion of less responsive chirality) and better overall photoluminescent intensity comparing from multi-chiral SWNT mixtures ^38^ [NO_PRINTED_FORM].

The reversibility of the IAA sensor was confirmed through monitoring its response to *in situ* photodegradation of IAA. IAA is photolabile and can be oxidized into OxIAA when exposed to low-wavelength high-intensity light ^39^. Our UV-Vis spectroscopy experiment showed a gradual degradation of IAA when exposed to a 600 nm light source (Fig. S6). Continuous monitoring of fluorescence intensity of the sensor showed that the addition of 5 μM IAA initially resulted in fluorescence quenching, but gradually recovered upon prolonged excitation, indicating in situ photodegradation of IAA to OxIAA (Fig. 2e). Similar observations were made upon repeated additions of 5 μM IAA to the same sample (Fig. 2e). This result indicates that the binding between IAA and its sensor is reversible, and the kinetic response is on a similar time scale as the rate of IAA photodegradation (Fig. S6)

### IAA nanosensor localized inside leaf cells upon infiltration

For accurate interpretation of the IAA nanosensor signal *in planta*, we need to determine its localization in the plant tissue. While IAA is biosynthesized in the cytoplasm, it may be transported to the nucleus for regulating auxin-responsive genes, conjugated and catabolized in the endoplasmic reticulum, or sequestered in the vacuole to regulate available IAA in plant cell ^40,41^ Furthermore, IAA also undergoes intercellular transport to establish auxin gradients along plant tissues ^42^. Although numerous transgenic biosensors have successfully demonstrated IAA detection inside roots, our study places its initial emphasis on IAA detection within leaf tissues, as IAA also contributes to leaf development^3^ [NO_PRINTED_FORM] but existing fluorescent biosensors are limited by chlorophyll autofluorescence in leaves ^24,25^. Therefore, we administered the SWNT-based IAA nanosensor into leaf tissues through the abaxial epidermis using a needleless syringe. Infiltration of nanosensors by this technique did not affect plant growth or physiology, as there was no visible damage to leaf tissues nor reduction of chlorophyll content in leaves infiltrated with the nanosensor (Fig. S7). The nanosensor is designed using its size and zeta potential to enable spontaneous transport through the cell wall and lipid membrane, as described in our LEEP theory^43^.

To investigate the localization of the IAA nanosensor after infiltration into leaf tissues, we used a confocal Raman spectroscopy system to precisely detect the SWNT-specific Raman G-band at the cellular level. *In vitro* Raman measurement of the IAA nanosensor verified that it produces a strong Raman G-band at 1590 cm^−1^ Raman shift (Fig. 3a). Using Arabidopsis leaves infiltrated with the IAA nanosensor, we measured the Raman spectra of distinct cells in the transverse cross-section and abaxial epidermal surface of the leaf (Fig. 3b). Interestingly, we found that the IAA nanosensor did not localize equally to the various cells of the leaf tissue. The IAA sensor Raman peak showed the highest intensity in the spongy mesophyll cells, followed by the palisade mesophyll cells (Fig. 3c). In contrast, the air space and epidermal cells, such as guard cells and pavement cell displayed relatively low sensor Raman peak intensity (Fig. 3c). This suggests that, although the IAA nanosensor was infiltrated through the abaxial leaf surface, the sensors selectively localized to the mesophyll cells.

**Fig. 3.**
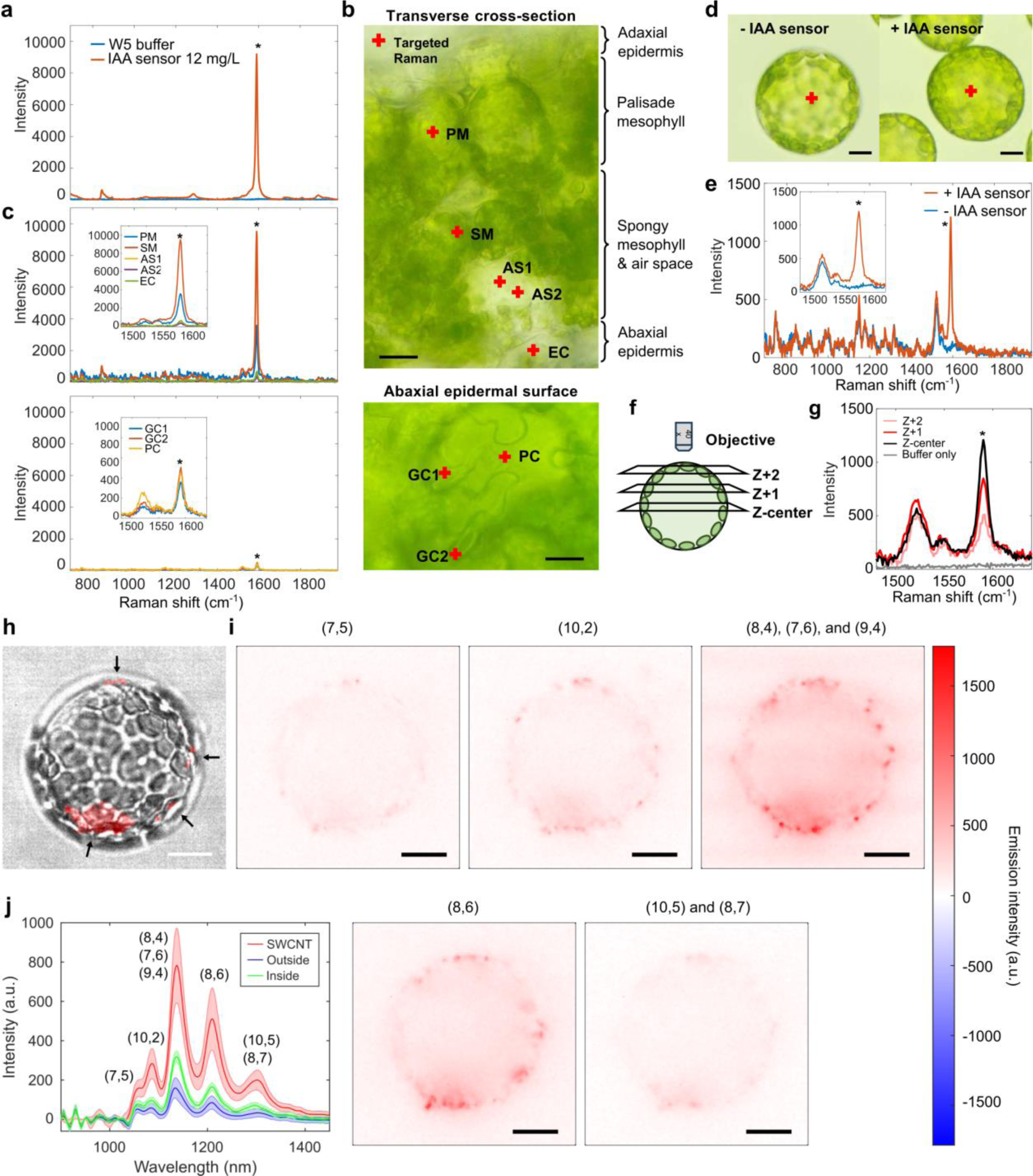
IAA nanosensor localizes into plant cells. **(a)** Raman spectra of *in vitro* SWNT-based IAA nanosensor and W5 buffer. The IAA nanosensor generates the distinctive 1590 cm^−1^ Raman G-band, while the W5 buffer does not contain any strong Raman signal in the detected spectra. **(b)** Bright field images of cells in Arabidopsis leaf blade after infiltration with IAA nanosensor. Top, transverse cross-section. Bottom, abaxial leaf epidermis. Locations indicated by a red cross were subjected to confocal Raman spectroscopy. PM, palisade mesophyll cell. SM, spongy mesophyll cell. AS, air space. EC, epidermal cell. GC, guard cell. PC, pavement cell. Scale, 20 µm. **(c)** Raman spectra of individual leaf cells and structure in **(b).** Inset, zoomed in Raman spectra showing 1590 cm^−1^ IAA nanosensor G-band. **(d)** Protoplasts isolated from Arabidopsis leaves with and without IAA nanosensor infiltration. Locations indicated by a red cross were subjected to confocal Raman spectroscopy. Scale, 10 µm. **(e)** Average Raman spectra of protoplasts in **d** (*n*=3). Inset, zoomed in Raman spectra showing 1525 cm^−1^ carotenoid peak and 1590 cm^−1^ IAA nanosensor G-band. **(f)** Schematic diagram of optical-sectioning focal planes from center (Z-center) to surface (Z+2) of the protoplast. **(g)** Raman spectra of the protoplast center at various focal planes, as illustrated in **e**. Inset, zoomed in Raman spectra showing 1525 cm^−1^ carotenoid peak and 1590 cm^−1^ IAA nanosensor G-band Asterisk (*) indicates IAA nanosensor G-band at 1590 cm^−1^ Raman shift. **(h)** Bright field image of isolated *N. benthamiana* protoplast incubated with 5 mg L^−1^ IAA nanosensor overlaid with sensor nIR fluorescence of the sensor (red) under 730 nm excitation. Black arrows indicate the localization of the sensor fluorescence signal inside the isolated protoplast. Scale bar, 5 μm. **(i)** Localization of IAA nanosensor of different chiralities in the periphery of the isolated protoplast. Similar distribution was observed in 12 biological replicates (*n*=12). Scale bar, 5 μm. **(j)** IAA nanosensor fluorescent spectra obtained from the pixels containing the IAA nanosensor (pixels with total fluorescent intensities integrated across 900 – 1450 nm greater than 6600), randomly selected pixels in the surrounding media outside the isolated protoplast, and randomly selected pixels in the center of the isolated protoplast. Data are mean ± s.d. from all the selected pixels.

To elucidate whether the IAA nanosensors entered the leaf cells or retained on their cell surface after infiltration, we isolated protoplasts from leaf tissues with and without IAA nanosensor infiltration. Protoplasts isolated from both samples were visually similar, suggesting that the IAA nanosensor did not adversely affect the cells (Fig. 3d). Confocal Raman spectroscopy at the center of these protoplasts revealed that the sensor-specific Raman peak was found only in protoplasts isolated from leaf tissue infiltrated with the IAA nanosensor (Fig. 3e). This implies that the IAA nanosensors localized inside the Arabidopsis leaf cells after infiltration. To further investigate if the IAA nanosensor specifically localized to the center of protoplasts, we performed Raman spectroscopy measurements along the plasma membrane, as well as a nearby area containing only the buffer (Fig. S8a). The sensor-specific Raman peak was detected only at the center of the protoplasts. In contrast, the Raman spectra along the plasma membrane (up, down, left, right, Fig. S8b) comprised of strong carotenoid Raman peaks that corresponded to the abundance of chloroplasts, but no detectable sensor Raman peak (Fig. S8b). Furthermore, we measured the Raman spectra at different focal planes to ascertain if the IAA nanosensor is found inside the protoplast (Fig. 3f). While the sensor-specific Raman peak showed the highest peak intensity at the center focal plane, shifting the focal plane towards the protoplast surface resulted in progressively weaker sensor Raman peak intensity (Fig. 3g). These suggest that the IAA nanosensor was found inside the protoplasts and not on the cell surface.

The IAA nanosensor localization inside leaf cells was further confirmed using *Nicotiana benthamiana*. Similar to Arabidopsis, the sensor Raman peak intensity was strongest in the spongy mesophyll cells of *N. benthamiana* leaf tissue (Fig. S9a-b). In contrast, other cell types and the air space displayed much weaker sensor Raman peaks (Fig. S9a-b). This implies that the IAA nanosensor mostly localized to the spongy mesophyll cells in *N. benthamiana*. Likewise, the sensor Raman peak was detected in protoplasts isolated from *N. benthamiana* leaf samples infiltrated with the IAA sensor (Fig. S9c-d). In contrast, leaves without sensor infiltration did not produce any protoplast with the sensor Raman peak (Fig. S9c-d). The highly similar results between Arabidopsis and *N. benthamiana* indicates that the IAA sensor localized inside leaf cells in most plant species upon infiltration.

The ability of the IAA nanosensor to localize within isolated *N. benthamiana* protoplasts was further supported by our hyperspectral study (Fig. 3h-j). After 1 hour incubation of the IAA sensor with the isolated protoplasts, we observed localization of the IAA nanosensor nIR fluorescence signals within the protoplasts, demonstrating passive transport across the cell membrane without external force (Fig. 3h). After deconvoluting the total fluorescence into each chirality, we observed that all sensor chiralities show similar uptake result where the sensor, indicated as the sharp red dots, localizes to the periphery of the protoplast (Fig. 3i). We further prove the identity of the sharp SWNT dots to be the IAA sensor through its signature SWNT fluorescence spectra as shown in Fig. 3j. The spectra of pixels in the buffer outside the protoplast and deep inside the protoplast are due to scattering of the SWNT emissions by the local media, causing non-SWNT locations to show weak fluorescence (Fig. 3j). These results suggest that the IAA nanosensor can be applicable across different plant species.

### IAA nanosensor allows for quantification of IAA levels in Arabidopsis plant

The results from the *in vitro* investigation showed that IAA nanosensor could exhibit dose-dependent quenching response to IAA. To validate and study the response of the sensor *in planta*, we developed a transgenic Arabidopsis plant expressing *XVE::iaaM*, which uses a β-estradiol inducible gene expression system to control endogenous IAA levels ^44^. *XVE::iaaM* Arabidopsis plants were screened and an optimal transgenic line was selected to have minimal basal expression but strong and proportional *iaaM* expression upon treatment with the β-estradiol inducer (Fig. S10). Topical application of β-estradiol on the leaf resulted in leaf curling and hyponasty, which are indicative of an increase in endogenous IAA and auxin activity (Fig. 4a, S11a-b). The induction of *iaaM* gene expression, auxin-responsive *GH3.3* gene expression, and endogenous IAA level at 24 h post-treatment were verified upon treatment with β-estradiol inducer (Fig. 4b).

**Fig. 4.**
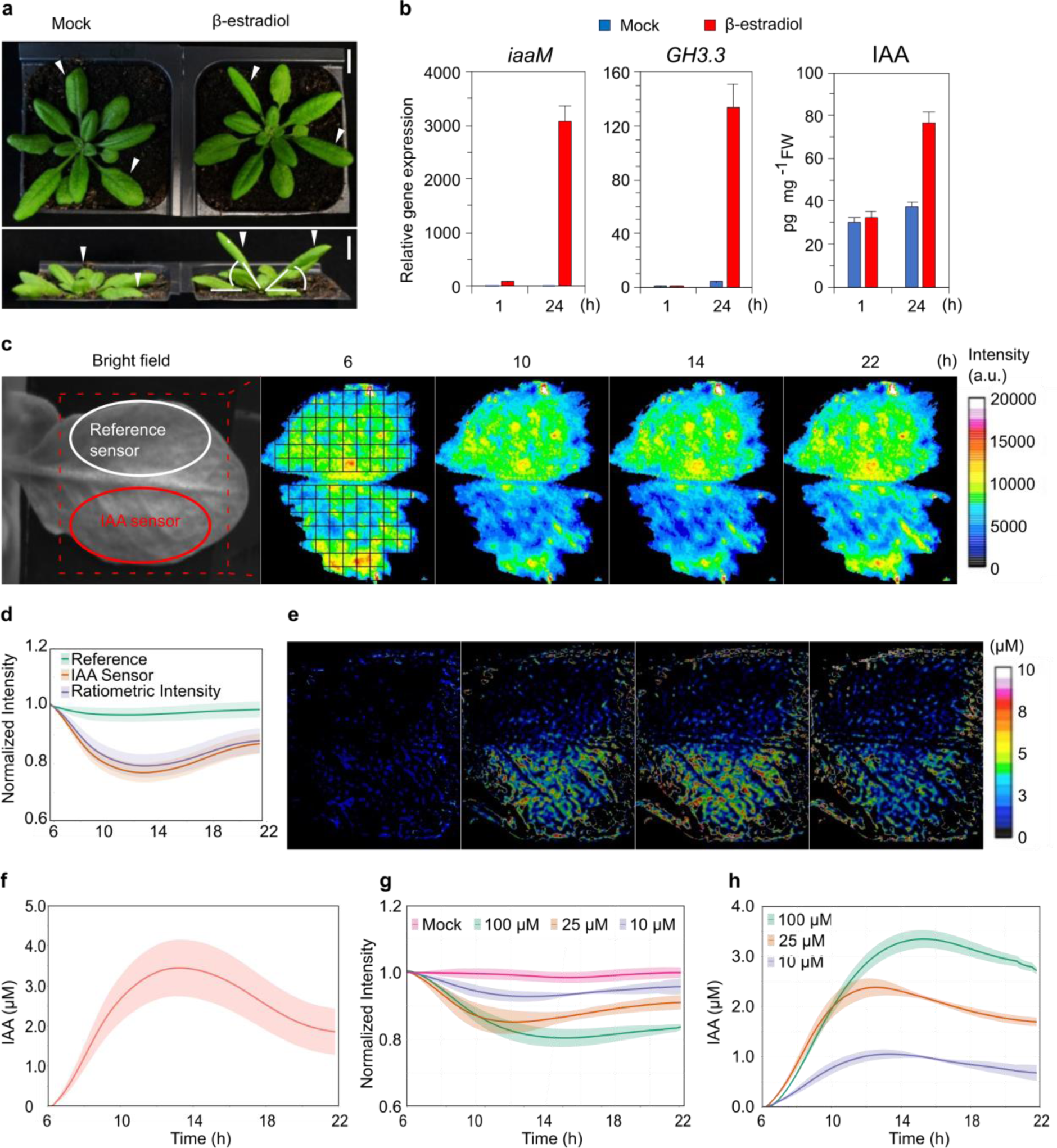
*In vivo* validation of IAA nanosensor in *XVE::iaaM* Arabidopsis plant. **(a)** Auxin-induced morphological changes in 4-week-old Arabidopsis plants expressing *XVE::iaaM*. Arrowheads indicate leaves treated with DMSO (mock) or 100 µM β-estradiol for 24 h. Scale bars: 5 mm. **(b)** Expression of *iaaM* and auxin-responsive gene GH3.3, and IAA level in *XVE::iaaM* Arabidopsis plants after 1 and 24 h of inducer or DMSO (mock) treatment. Data are mean ± s.d. from three biological replicates (*n*=3). **(c)** nIR fluorescent images showing intensities of reference sensor (upper half) and IAA sensor (bottom half) at different time points. Left: Bright field image of Arabidopsis rosette leaf used for application of reference sensor (white oval) and IAA sensor (red oval) using needle-less syringe infiltration. After topical application of the inducer or mock treatment, plant samples were allowed to rest for 6 h before nIR fluorescence imaging. Three independent experiments were repeated with similar results (*n*=3). **(d)** Time-profile average normalized intensity (I/I_o_) of IAA sensor, reference sensor, and ratiometric intensity obtained from **c**. Io represents the average fluorescent intensity measured during the first 5 minutes of the experiment. Corresponding shades represent the standard deviation from each ROI (black squares). **(e)** IAA distribution at each corresponding time point in **c** estimated using our correlation model between ratiometric signal and local IAA concentration (see supplementary information). **(f)** Time-profile average IAA concentration obtained from **d** and **e**. **(g)** Average normalized ratiometric responses to different β-estradiol concentrations. Mock: DMSO treatment. Corresponding shades represent the standard deviation from the mean of three independent biological samples (*n*=3). **(h)** Corresponding average IAA concentration profiles estimated from **h**.

While the techniques described above elucidated IAA content and the corresponding auxin activity at specific time points, use of the IAA nanosensor *in planta* would enable real-time analysis of auxin dynamics upon treatment with β-estradiol. In this work, we utilized a ratiometric platform ^28,38^ consisting of IAA nanosensor and an inert reference nanosensor, (AT)_15_-SWNT, for *in vivo* detection of endogenous IAA (Fig. 4c). Our previous work demonstrates how such nanosensors can be utilized ratiometrically ^38^. (AT)_15_-SWNT is a nano-conjugate formed from the wrapping of single-strand DNA (AT)_15_ on the nanotube surface. This inert sensor gives invariant optical response to IAA and other relevant plant analytes (Fig. S12). The use of a reference sensor improves the confidence of attributing the observed response of IAA nanosensor to the presence of IAA in a complex *in vivo* medium. In addition, the ratiometric response, calculated as the ratio of IAA nanosensor fluorescence intensity to the average intensity of reference sensor, would improve the signal-to-noise ratio and provide a more accurate evaluation of IAA concentration. This is because any fluctuation in fluorescence signal due to external factors, such as laser power fluctuations or background lighting, would be reflected by the reference sensor response, which can then be corrected by ratiometric analysis.

By using a needleless syringe, the IAA nanosensor and the reference nanosensor were infiltrated into two adjacent regions of the rosette leaf lamina that were separated by the midrib (Fig. 4c). The nIR fluorescence of both nanosensors was monitored under constant excitation at a standoff distance of 1 m using a 2D array InGaAs detector equipped with a 900 nm long-pass filter. The use of a standoff detection setup enables remote sensing with an expanded field of view, facilitating the study of whole-plant physiology with minimal sample handling. After 6 h topical treatment of 100 μM β-estradiol on the adaxial surface, fluorescence intensity of the IAA nanosensor and reference sensor were captured continuously for 16 h. False-color nIR images of the leaf sample at different time points showed increasingly potent systematic fluorescent quenching of the IAA nanosensor, from 6 to 10 h post-treatment of β-estradiol, followed by a gradual return towards the baseline from 10 h onwards (Fig. 4c-d). On the other hand, the fluorescence intensity of reference sensor (AT)_15_-SWNT remained relatively constant throughout the experiment (Fig. 4c-d). The significant quenching observed by the IAA nanosensor indicated an increase of IAA content (Fig. 4d), which is aligned with the result from biochemical assays (Fig. 4b). To quantitatively assess the change in IAA levels induced by β-estradiol, we correlated the ratiometric response of IAA nanosensor to IAA concentration using a reversible first-order binding model for the IAA nanosensor (see Supplementary Information for model development). In this model, we utilized the normalized fluorescence intensity I/I_o_ which was calculated by using the average intensity of the first 100 scans as the baseline (I_o_). As such, the derived result should be interpreted as the change of auxin level compared to the initial time point before induction. By extracting the ratiometric response from each pixel and applying it to the model, we generated 2D images of auxin distribution level from the corresponding original nIR images (Fig. 4e). These images showed a substantial increase in IAA level from 6 to 22 h after treatment, with the increase being uniform across the lamina separated by the secondary veins (Fig. 4e). In contrast, no spatial distribution of auxin levels was observed in the upper half area of the leaf where the reference sensor was applied (Fig. 4e). Fig. 4d and 4f summarize the average time-profile response of the reference nanosensor, the IAA nanosensor, its ratiometric value, and corresponding calculated changes in IAA level; showing that chemically induced IAA biosynthesis peaks at around 13 h after treatment, followed by a gradual decrease. The reduction in IAA concentration after the peak could be attributed to IAA metabolism in response to excess IAA production, as evidenced by the increased expression of *GH3.3* (Fig. 4b).

The nanosensor also demonstrated a dose-dependent response to β-estradiol. *XVE::iaaM* Arabidopsis leaves treated with 0 µM (mock), 10, 25, and 100 µM of β-estradiol resulted in the IAA nanosensor to show transient modulations in intensity, which were relatively proportional to the amount of β-estradiol applied (Fig. 4g). The mock treatment resulted in a constant ratiometric intensity, while higher concentrations of β-estradiol caused greater signal quenching (Fig. 4g-h). This highlights the sensitivity of our IAA nanosensor in measuring transient changes to IAA levels.

### Detection of local IAA movement in *XVE::iaaM* Arabidopsis plants using IAA sensor

Spatiotemporal distribution of auxin is a key factor in regulating plant growth and development ^12,45^. By using *XVE::iaaM* Arabidopsis plants, the IAA nanosensor is able to track the transport of chemically-induced IAA. While the previous experiment applied the β-estradiol inducer to the entire leaf to cause a global increase in IAA levels, topical application of the inducer at a small, specific spot on the leaf would give rise to a localized increase in IAA levels, which is useful for demonstrating auxin movement. Spot application of the inducer on one side of the leaf triggered leaf curling only on the treated side and hyponasty, indicating a localized increase in IAA concentrations (Fig. 5a, S12c). Similar to the previous experiment (Fig. 4b), the localized induction of *iaaM* and subsequent IAA accumulation was verified by gene expression of *iaaM* and *GH3.3* (Fig. 4c). *iaaM* expression was greatly induced at the treated spot, while expression in the treated side and the untreated side of the leaf remained low (Fig. 5a,c), indicating that the inducer did not diffuse away from the treated spot and that inducer-enhanced IAA biosynthesis occurred only at the treated spot. However, the expression of auxin-responsive gene *GH3.3* was strongly induced at the treated spot compared to the untreated side (132-fold increase), while the treated side also showed a significant but milder induction (31-fold increase) and the untreated side had low *GH3.3* expression (Fig. 5c). This suggests that the induced local IAA synthesis at the treated spot was likely transported out to surrounding areas in the treated side of the leaf, but barely passed through the mid-vein to the untreated side. Likewise, application of the IAA nanosensor revealed a significant fluorescence quenching focused on the treated spot, which gradually expanded to the treated side over time (Fig. 5d-e). Image analysis showed a rapid increase in IAA level at the treated spot from 6 h post-treatment, which peaked at 14 h post-treatment, followed by a gradual decrease from 14 to 18 h post-treatment (Fig. 5d-f). In contrast, IAA levels in the surrounding areas on the treated side displayed a gradual increase, while the untreated side exhibited no significant change (Fig. 5d-f). Average time-profile curves were generated to visualize the IAA levels at the treated spot (T) and surrounding regions of interest (A, B, C, D; Fig. 5b, f), which clearly demonstrated IAA concentrations gradually increasing in the surrounding areas from 6 to 12 h after treatment. This indicates that IAA biosynthesis was induced at the treated spot, then transported to the surrounding cells and tissues. At the same time, IAA level was unchanged on the untreated side (U, Fig. 4f), which is consistent with auxin-responsive gene expression (Fig. 5c) and a previous study that showed the midvein acting as a boundary for auxin response ^46^.

**Fig. 5.**
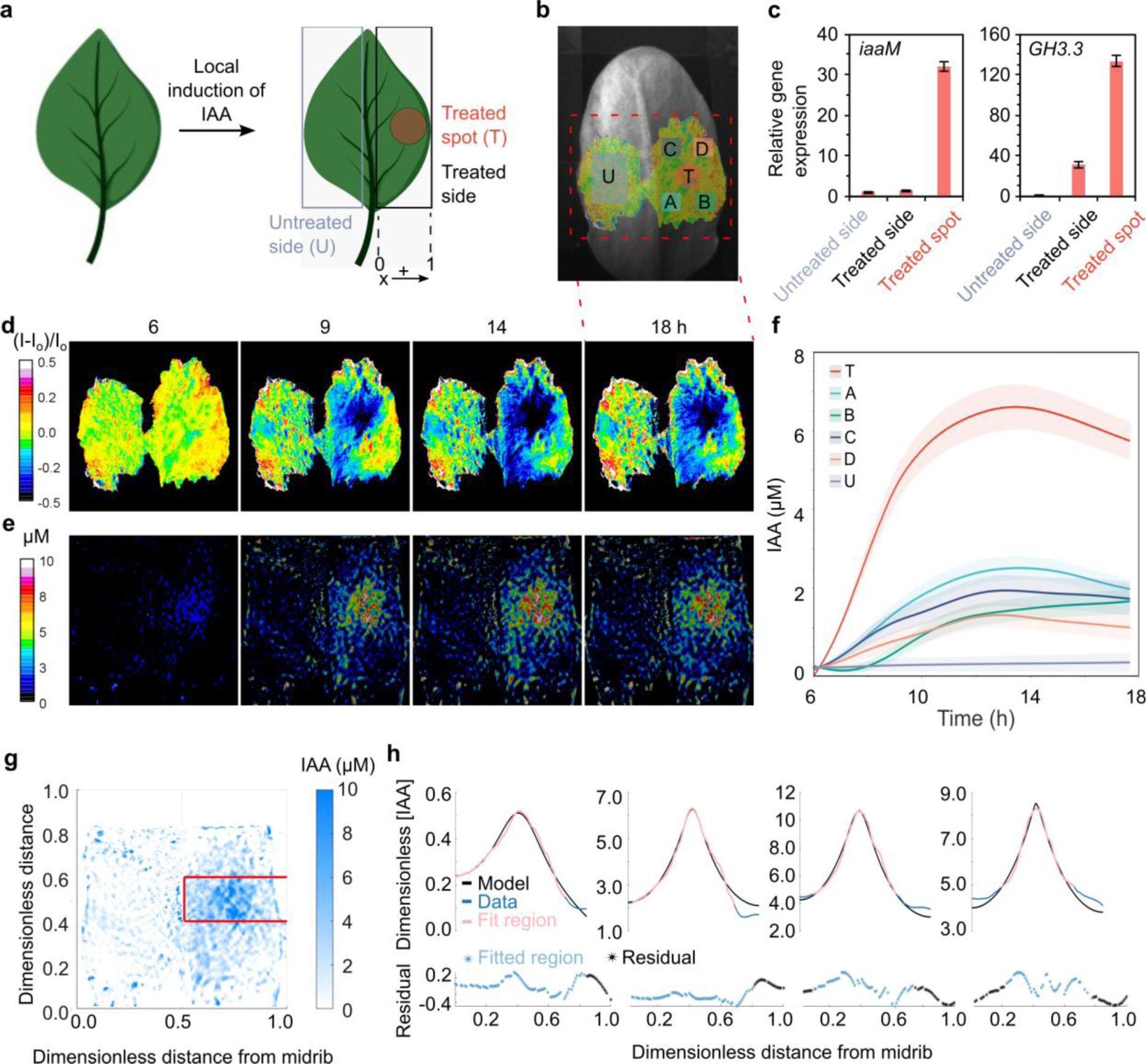
Local Transport of Auxin in Arabidopsis Leaf. **(a)** Schematic diagram illustrating β-estradiol treatment spot and other parts of the treated leaf. β-estradiol was locally applied in spherical treated spots near the leaf margin on one side of rosette leaf blades of *XVE::iaaM* Arabidopsis plants. The model domain for auxin transport analysis is a one-dimensional line on the β-estradiol-treated half of the leaf, spanning from the midvein to the leaf margin. **(b)** Bright field image of a typical Arabidopsis rosette used for local transport of auxin experiment. ROIs A, B, C, D, T, and U are colored on top of the image. U: untreated side; T: treated spot; A: toward base spot at midvein side; B: toward base spot at edge side; C: toward apex spot at midvein side; D: toward apex spot at edge side. **(c)** *iaaM* and GH3.3 gene expressions in β-estradiol-treated spot, treated side (excluding treated spot), and untreated side. Sample collected from the treated side excludes the treated spot. Data are mean ± s.d. from three biological replicates (*n*=3). **(d)** Normalized fluorescent intensity images of leaf sample in **b** at different time points showing quenching response of IAA sensor at treated area. After topical application of the inducer at the treated spot, plant samples were allowed to rest for 6 h before nIR fluorescence imaging. **(e)** Estimated IAA distribution of leaf sample in B at different time points. **(f)** Average time-profile concentration curves of IAA at designated spots on treated Arabidopsis leaf. Shaded regions indicate standard deviation of pixels in selected regions of interest. **(g)** A strip of data centered around the induced spot was averaged into one dimension for data fitting with model. **(h)** Dimensionless model (Equations 2-4) was fit to data at each time point shown above using parameters σ ̅, γ ̅, ϕ, and k_margin_. Only light magenta region labeled “Fitted Region” was used for fitting model to avoid artifacts at edges of sensor spots. A residual plot of [*IAA*]_*model*_− [*IAA*]_*data*_is shown beneath each fit. Blue points in residual plots correspond to “Fitted Region” points.

To further probe the spatial distribution and transport of IAA, we developed a one-dimensional steady-state reaction-diffusion model to describe the spatial IAA concentration data in Fig. 4C-E at each time point. Specifically, the concentration profile from the midvein (*x* = 0) to the margin of the leaf (*x* = 1) was analyzed (Fig. 4g). In our model, IAA is generated with rate *γ̅* across the inducer-treated spot with its distribution modeled by a Gaussian function centered about point *a̅* with standard deviation *σ̅*. IAA is free to diffuse within the leaf but is also inactivated and consumed throughout the leaf. We define a Damköhler number *ϕ*^2^ to describe the relative rates of IAA reaction and diffusion. At the midvein, we employ a no-flux boundary condition, as IAA was shown to be unable to cross the midvein by our data (Fig. 4c-f) and a previous study ^46^. As for the leaf margin boundary, we considered that auxin response has previously been shown to accumulate at the leaf margins under shade conditions ^46^. It has been suggested that auxin transport proteins are localized along the cells to form a canal of auxin flow from source to the sink ^47^. We thus employ a convective boundary condition between the IAA concentration in the leaf blade and the bulk IAA concentration in the leaf margins with mass transfer coefficient *k*_*margin*_. The non-dimensionalized mass balance for IAA can be written as follows (see Supplementary Information for details):

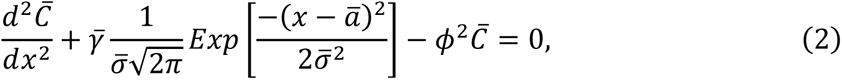

with boundary conditions

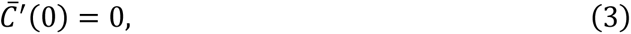

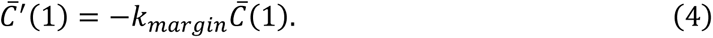

*C̅* is the dimensionless concentration of IAA and *x* is a dimensionless length scale. For fitting to the data, we chose to analyze a small region approximately centered about the induced spot as a representative region (Fig. 5g). Data close to the edges of the sensor spot were neglected due to artifacts. The model parameters *σ̅*, *γ̅*, *ϕ*, and *k*_*margin*_ were fit to the experimental data at each time point (Fig. 5h). The results for each highlighted time point in Fig. 5g are summarized in Table 1. *σ̅* and *γ̅* are shape parameters reflecting the production magnitude and spread of IAA around the inducer-treated spot.

**Table 1.** Model Fitting Parameters.

|  | 6 h | 9 h | 14 h | 18 h |
| --- | --- | --- | --- | --- |
| $\bar{\sigma}$ | 8.3E-02 | 3.3E-02 | 4.3E-02 | 1.2E-02 |
| $\bar{\gamma}$ | 4.2 | 47 | 86 | 45 |
| $\phi$ | 3.4 | 3.6 | 3.8 | 2.9 |
| $k_{margin}$ | 9.2E+00 | 5.4E-03 | 3.9E-02 | 6.0E-05 |

Interestingly, *ϕ*^2^ appears to be relatively consistent across the different time points. Physically, *ϕ*^2^ ≫ 1 corresponds to IAA that is rapidly consumed and cannot diffuse far across the leaf, whereas *ϕ*^2^ ≪ 1 corresponds to IAA that is free to travel far across the leaf without being reacted away. Here, we find *ϕ* > 1, which corresponds well with the data in Fig. 5d-f. IAA slowly spreads across the leaf, reflected by the gradually increasing concentration profiles at the analyzed spots (A, B, C, D; Fig. 5b, 5f). However, the IAA concentrations at the analyzed spots never equalize with the concentration at the treated spot (T; Fig. 5b, f), indicating rapid degradation. In our model, the Damköhler number is a ratio of two length scales: 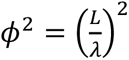, where *λ* ≅ 0.14 cm is the length of the reaction zone (see Supplementary Information for details). According to the model, IAA can diffuse slightly under one-third the length of the treated half of the leaf before it is fully consumed. *k*_*margin*_is highest at the first time point (6 h), but then drops off in the later time points. This suggests that although the margin concentration may be negligible initially, it may begin to “fill” with IAA, reducing the driving force for mass transfer. Notably, the model is insensitive to *k*_*margin*_at the later time points (*t* ≥ 9 h), suggesting that the IAA concentration in the margin may be saturated by the IAA produced in the blade of the leaf, resulting in a boundary condition resembling no flux, like the midvein boundary condition. This can be seen in the behavior of the model solutions for later time points (Fig. 5h, *t* ≥ 9). This analysis is able to describe the IAA concentration maximum as between the mid-rib and leaf margin, and therefore supports the observation of a no flux boundary condition at the former location.

## Discussion

Auxin IAA is a vital hormone that contributes to plant development and its response to environmental stresses. Therefore, accurate quantification and visualization of its levels *in vivo* are critical to understand its function in plant research ^48^. However, existing IAA measurement methods have limitations. Indirect methods using gene reporter systems, such as *DR5::GUS* and *DII-VENUS*, have potential problems of cross-reactivity issues with other plant hormones^49–51^, inability to detect transient changes ^15^, and species- and tissue-dependent responses ^52,53^. However, quantification of *in vivo* IAA concentration still requires the use of LC-MS, which requires homogenization and lacks spatial resolution ^54,55^. This issue is addressed by the recent development of a FRET-based IAA biosensor, which allows for direct and specific detection of IAA *in vivo* ^16^. Nevertheless, all these genetically encoded biosensors require genetic engineering of target plants and are primarily used for imaging of the root system rather than green tissues like cotyledons and leaves ^16^. While IAA also plays an important role in regulating leaf growth and development, including cell differentiation, patterning, morphogenesis, and response to environmental stimuli ^56^, detecting and visualizing IAA distribution in leaf tissue can be challenging due to its complex structure, as well as the presence of pigments and waxes that can interfere with the detection of IAA. Additionally, as the distribution of IAA is spatially heterogeneous, high accuracy and spatial resolution when measuring IAA concentrations *in planta* are useful to elucidate its effects on leaf development. However, some conventional methods may not adequately resolve such differences.

To address this challenge, we leverage on plant nanobionics technology, an emerging field that intersects plant biology and nanotechnology, to develop a new approach for direct visualization of auxin *in planta*. By using nanomaterials, novel sensors, and interfaces can be designed to detect a wide range of analytes ^27,48^, which can be customized for specific applications with greater sensitivity and selectivity. For instance, we have successfully developed a library of biocompatible nanosensors that can detect and decode endogenous H_2_O_2_ signals in plant stresses ^28^, trace exogenous synthetic auxin *in planta* ^57^, and monitor arsenic uptake in the root system^29^.

In this study, we present a novel CoPhMoRe-based IAA nanosensor that allows for real-time spatio-temporal imaging of IAA in living plants. Specifically, our sensor has the following advantages: (i) biomimetic design to resemble the natural binding site between IAA and TIR1, which confers selectivity and sensitivity to IAA *in vitro* and *in viv*o, (ii) direct and reversible binding of IAA for accurate measurement of IAA concentrations and the ability to detect transient changes, (iii) high stability of sensor and fluorescence signal, which allows for reliable time-course experiments, (iv) enables visualization of IAA distribution and movement in live leaf tissue, (v) easy and fast application across plant species without any genetic engineering or optimization for sensor application. For validation of the IAA nanosensor, we firstly used *XVE::iaaM* Arabidopsis plants, which has controllable endogenous IAA levels that depend on the application of an external inducer. Importantly, our results showed that the nanosensor exhibited a dose-dependent response to varying IAA levels, which corresponded to the concentration of the inducer treatment. Furthermore, our results demonstrated the nanosensor’s capability in tracking local IAA content and transport via 2D mapping of the fluorescent intensities. A one-dimensional steady-state reaction-diffusion model was developed to analyze the spatial distribution and transport of IAA in a leaf treated with single spot of inducer, with a no-flux boundary condition at the midvein and a convective boundary condition at the leaf margin. By fitting with the experimental data, our model suggests that IAA synthesized at the treated spot undergoes gradual diffusion to surrounding parts of the leaf, as IAA moves to almost one-third length on the treated half of the leaf before it is fully consumed or metabolized. While the genetically encoded biosensors provide excellent visualization of auxin distribution, they lack quantitative measurement of auxin levels. In contrast, the IAA nanosensor provides both qualitative and quantitative data, but the performance of the sensor may be affected by the complex *in vivo* medium. Nevertheless, the dose-dependent response of the IAA nanosensor to varying IAA levels, as controlled by the inducer in *XVE::iaaM* Arabidopsis plants, provides valuable insight into the sensor’s performance and can be used for *in vivo* calibration. However, there are still some drawbacks that need to be addressed to improve the sensor’s performance. For instance, the current method of analyzing the sensor’s response does not provide information on the absolute basal auxin level at the time of measurement, which is critical for accurate quantification. To solve this issue, an internal reference sensor can be incorporated, either by using single chirality SWNTs for selective fluorescence with specific excitation wavelength ^38^ or by combining the spectrofluorometer with Raman spectroscopy to use the characteristic G-band as an internal reference ^58^. These methods can provide accurate scaling of the nIR fluorescence intensity without the need for an external reference sensor. The current stand-off detection set up allows spatial resolution of the whole leaf but may not provide detailed cellular-level information. It still serves a valuable purpose in capturing the overall auxin distribution and dynamics at a macroscopic level. The spatial resolution of the whole leaf is useful in providing an overview of auxin accumulation patterns across different regions of the leaf, identifying spatial gradients, and detecting broader trends in auxin distribution. This information can be crucial for understanding the global regulation of auxin and its impact on development ^56^. While this study focused on the performance and testing of the IAA nanosensor on leaf tissue, it has the potential to be used in other plant tissues with appropriate delivery techniques such as microneedles-assisted target delivery ^59^, pressurized chamber ^60^ or vacuum infiltration ^61^ for whole plant infiltration. The IAA sensor introduced in this work may find utility in wide variety of studies and model systems, including investigations of the Arabidopsis root, apical hook, hypocotyl, developing leaf vasculature, shoot apical meristem. We have recently demonstrated the first use of such nanoparticle sensors in root systems ^61^. The IAA sensor is designed to cross cell wall and lipid membranes, as demonstrated in Figure 3, and therefore can be applied to a variety of cell and tissue types.

In conclusion, we have successfully developed a CoPhMoRe-based IAA nanosensor that enables direct visualization of IAA in living plants. Our sensor has distinct advantages over existing methods, including high stability, reversible binding of IAA, and the ability to detect transient changes. Using this sensor, we accurately measured IAA concentrations and visualized its distribution and movement in live leaf tissue under various conditions. Our work highlights the potential of plant nanobionics technology to address the challenge of measuring and visualizing auxin *in planta*. The development of this IAA nanosensor provides a new tool for plant researchers to investigate the role of IAA in plant growth and development. Moreover, it can be utilized as a diagnostic tool for early detection of plant stress in agriculture. Looking ahead, we envision that our sensor technology could be further explored for the detection of other plant hormones and signals, presenting a powerful tool for plant biology and agriculture research.

## Methods

### General procedure for synthesis of polyamic sodium salts

All reagents were purchased from Sigma Aldrich and Tokyo Chemical Industry (TCI). A typical procedure for the synthesis of polyamic sodium salt is as follows. In a round bottom flask under argon atmosphere was added one molar equivalence of dianhydride and one molar equivalence of diamine, followed by addition of anhydrous N-methylpyrrolidone (NMP) (10 wt%). After stirring at room temperature for 24 h, excess volume of NaOH 0.1N was added dropwise to the reaction mixture to completely neutralize the carboxylic acid groups. Precipitation was initially observed which gradually dissolved to give a clear solution with pH ∼13 - 14. The resulting solution was dialyzed against deionized (DI) water using regenerated cellulose membrane (12-14 kDa molecular weight cut-off) for 48 h where the water reservoir was refreshed every 12 h. After dialysis, the aqueous solution was frozen, followed by freeze-drying to obtain the polymer as a dry powder. The polymers were stored at −20 °C for later experiments.

### General procedure for preparation of polymer-SWNT complexes (1)−(20)

A suspension of 5 mg purified high-pressure carbon monoxide SWNTs (NanoIntegris, lot # HP32-018) in corresponding aqueous polymer solution (25 mg, 5 mL DI water) was tip-sonicated at 22% amplitude (Qsonica, pulse mode: 5s ON, 2s OFF) for 1 h in an ice-bath. The resulting suspension was then ultra-centrifuged (35,500 rpm) for 4 h. Approximately 80% of the upper supernatant was carefully collected. SWNT concentration was estimated by UV-Vis spectrophotometry (Agilent Cary 5000) using SWNT absorbance at 632 nm with an extinction coefficient of 0.036 (mg L^−1^)^−1^ cm^−1^.

### *In vitro* screening of plant hormone analytes

For plant hormone screening, 1 mL of stock solutions containing 100 mM of each analyte dissolved in dimethyl sulfoxide (DMSO) for indole-3-acetic acid (IAA), indole-3-butyric acid (IBA), abscisic acid (ABA), gibberellic acid (GA_3_), salicylic acid (SA), and methyl jasmonate (MeJA), or in NaOH 100 mM for IAA were prepared. Stock solutions of SWNT suspension with a concentration of 2 mg L^−1^ in 2-(*N*-morpholino)ethanesulfonic acid (MES) buffer (10 mM, pH 5.5) were also prepared. SWNT fluorescence was measured in a quartz cuvette (1 μL of analyte solution was added to 999 μL of SWNT solution in a quartz cuvette) using home-built stand-off nIR camera set up with excitation wavelength of 785 nm and laser power of 30 mW. Fluorescent intensity was measured before and after the addition of analytes.

### Confocal Raman spectroscopy of leaf tissue and isolated protoplasts

To isolate protoplasts, leaves were cut into strips of 1 mm width, which were then submerged in the digestion buffer [1% (w/v) Cellulase Onozuka R-10 (Yakult Pharmaceutical Industry, Japan), 0.25% (w/v) Macerozyme R-10 (Yakult Pharmaceutical Industry, Japan), 0.4 M mannitol, 20 mM MES (pH 5.7), 20 mM KCl, 10 mM CaCl_2_]. The samples were subjected to vacuum infiltration for 10 min and subsequently placed in darkness at room temperature for up to 6 h. Afterwards, the samples were filtered using Miracloth (Calbiochem, USA) to obtain the protoplast suspension. The protoplasts were allowed to sink to the base of the tube before the supernatant was discarded. Subsequently, equal volume of W5 buffer [154 mM NaCl, 125 mM CaCl_2_, 5 mM KCl, 2 mM MES (pH 5.7)] was added to resuspend the protoplasts. The protoplasts were washed twice more in a similar manner, then resuspended in W5 buffer for confocal Raman spectroscopy.

Leaf tissue and isolated protoplasts were subjected to confocal Raman spectroscopy. A 40x magnification objective was used. Excitation wavelength of 785 nm was used. A total of 5 spectra was measured per location, with integration time of 2 s per spectra. Processing of Raman spectra were similar as previously described ^62^. Raman shift was calibrated against polystyrene, which has a well-documented Raman spectrum ^63^.

### Generation of transgenic Arabidopsis plants expressing *iaaM*

Pseudomonas *iaaM* gene (EC 1.13.12.3) was synthesized and cloned into pUC57 carrier vector (GenScript Biotech). The *iaaM* gene was then amplified with a Gateway adapter-flanked gene-specific primer pair (Supplementary Table 1), using pUC57-*iaaM* as the template, and cloned into Gateway® donor vector, pDONR221. The *iaaM* gene was subsequently cloned into pER8-DC, which contains an estrogen receptor-based transactivator (XVE) system^46^, through Gateway® recombination with pDONR-*iaaM* for inducible expression. The *iaaM* gene was also cloned into pBA-DC harbouring the constitutive promoter of cauliflower mosaic virus 35S (CaMV 35S) for transient expression in *N. benthamiana*. pER8-*iaaM* and pBA-*iaaM* plasmids were transformed into the *A. tumefaciens* strain, GV3101. Wild-type Arabidopsis (Col-0) was transformed with pER8-*iaaM* using the floral dipping method.

For the screening of *iaaM* expression in *XVE::iaaM* Arabidopsis lines, 10-d-old seedlings were transferred into ½ strength Murashige and Skoog (½ MS) liquid media with addition of 50 µM β-estradiol or DMSO as a mock-treatment control. The seedlings were then incubated overnight (16 h) under continuous white light (WL, Photosynthetic Photon Flux Density, PPFD = 100 µmol cm^−2^ s^−1^) with low agitation at 70 rpm. For concentration-dependent induction experiment, all growth conditions were the kept the same, but induction was carried out for 6 h using different inducer concentrations (0, 1, 10, 25, and 100 µM β-estradiol). For the time-dependent induction experiment, all growth conditions were the kept the same, but induction was carried out using 50 µM β-estradiol at various induction times (0, 1, 2, 6 and 14 h).

### *In vivo* detection of IAA nanosensor using home-build stand-off nIR camera system

In a typical experimental setup, a dilute aqueous suspension of 10 mg L^−1^ IAA nanosensor in MES buffer, corresponding to 15 ppm carbon, and aqueous suspension of reference sensor (single-stranded DNA (AT)_15_ wrapped CoMoCat (6,5)) were infiltrated onto the abaxial side of *Arabidopsis* or *N. benthamiana* leaves by a needleless syringe (Becton Dickinson). The infiltrated or nanosensor-functionalized leaves were allowed to rest for at least 1 h under growing condition before transferring to the home-built stand-off camera setup for nIR imaging. In particular, the nanosensor-infiltrated leaves were mounted and fixed vertically on the platform facing the camera, with a distance of 1 m. Two white LED light sources were also positioned alongside the plant to ensure it is not under any light stress. nIR fluorescence of the infiltrated leaf was imaged using a 785 nm laser with incident power of 20 mW. Each nIR image was captured with using a 2D array InGaAs detector equipped with a 900 nm long-pass filter, with integration time of 30 s and collected by Lightfield® software (Princeton Instruments).

For IAA detection in *XVE::iaaM* Arabidopsis plants, 100 µM β-estradiol supplemented with 0.01% Silwet L-77 solution was gently applied (on the whole leaf or a designated local spot) on the adaxial surface of 4-w-old Arabidopsis rosette leaf using a paint brush. After 2 h, IAA and reference nanosensor suspensions were infiltrated on abaxial side of the Arabidopsis rosette leaf. The plant was then placed in a growth chamber for another 4 h prior to nIR imaging experiment.

### Image analysis

Image and data analysis were primarily done using FIJI (Fiji Is Just ImageJ) and MATLAB 2021b. The raw image data obtained from the experiments were first deconvoluted with point spread function using our own home-built functions to restore and estimate the true fluorescent intensity of nanosensors without photon diffusion. To obtain a time profile of the nanosensor response, the pixel intensity of IAA and reference nanosensors was normalized to the corresponding initial values (first 5 minutes). The normalized IAA nanosensor intensity was then divided with normalized reference sensor intensity profile to obtain a ratiometric sensor response. Images of normalized intensity changes (I-I_o_)/I_o_ to display the false-colored sensor responses as in Fig. 3c, Fig. 4d (upper images), Fig. 5a and Fig. 5e were constructed by subtracting the first image from subsequent images then dividing the resulting images with the first image. Images showing IAA distribution in Fig. 3d, Fig. 4d (lower images), Fig. 5b and Fig. 5f estimated by the sensors were constructed by translating sensor response in each pixel to its corresponding auxin concentration using proposed auxin sensor binding model (See supplementary information).

### Quantification of free IAA

Samples from Arabidopsis and *N. benthamiana* were harvested and weighed, before being immediately frozen using liquid nitrogen. IAA phytohormone was then extracted using 80% methanol as described ^47^. Concentration of extracts was normalized to the fresh weight measured after harvest. IAA in the extracts was then quantified by Ultra Performance Liquid Chromatography−Tandem Mass Spectrometer (UPLC-MS/MS) equipped with UltiMate 3000 Rapid Separation LC system (Thermo Fisher Scientific) and the Q Exactive™ Hybrid Quadrupole-Orbitrap™ MS (Thermo Fisher Scientific). Accucore™ RP-MS column (Thermo Fisher Scientific) was used for compound separation. For the detection of IAA precursor ions, parallel reaction monitoring, targeted quantitation, and screening scan mode were conducted. 5 mM acetic acid in 5% (v/v) acetonitrile and in 95% (v/v) acetonitrile were used as mobile phases A and B, respectively. The elution profile was: 0–3 min, 5% mobile phase B in mobile phase A; 3–6 min, 5–95% mobile phase B in mobile phase A; 6–10 min 95% mobile phase B in mobile phase A; 10–10.1 min 95–5% mobile phase B in mobile phase A; and 10.1–11min 5% mobile phase B in mobile phase A. The mobile phase flow rate was 0.3 mL min^−1^. Injection volume was 5 µL. Electrospray ionization was operated in negative ion mode. TraceFinder™ 4.1 (Thermo Fisher Scientific) and Compound Discoverer 3.0 software (Thermo Fisher Scientific) were used to qualify the hormone levels. Hormone measurements were conducted on three biological replicates.

### RNA extraction and quantitative reverse transcription polymerase chain reaction (qRT-PCR)

Total RNA from plant samples was isolated using the GeneAll Ribospin RNA isolation kit (GeneAll). For cDNA synthesis, 1 µg of total RNA was reverse-transcribed at 42 °C for 90 min and 72 °C for 15 min using the moloney murine leukemia virus (M-MLV) reverse transcriptase (Promega). Each cDNA sample was diluted 8-folds and used for qRT-PCR with the specific primer pairs of each gene listed in Supplementary Table 1. qRT-PCR was performed using Takara SYBR Premix Ex Taq (Takara Bio) on Biorad CFX connect real-time system (Bio-Rad). Each reverse transcript was quantified in triplicate. The thermal cycling program was as follows: pre-denaturation at 95 °C for 2 min, denaturation at 95 °C for 10 s, followed by annealing and extension at 60 °C for 30 s, with a total of 40 cycles. The comparative threshold (Ct) cycle method (2–ΔΔCt) was used for relative quantification and calculations.

### Data availability

All data are provided in the article and the Supplementary Information or available from the corresponding authors upon reasonable request.

## Acknowledgements

This research was supported by the National Research Foundation (NRF), Prime Minister’s Office, Singapore under its Campus for Research Excellence and Technological Enterprise (CREATE) program. The Disruptive & Sustainable Technology for Agricultural Precision (DiSTAP) is an interdisciplinary research group of the Singapore MIT Alliance for Research and Technology (SMART) Centre.

## Author contributions

N.H.C., M.S.S., and I.C.J. conceived the project. M.S.S. and I.C.J. supervised the overall study. M.B.C.P., M.S.S., and I.C.J. designed the work. D.T.K., K.V.V. and B.J.R.S performed *in vitro* and *in planta* experiments and analyzed data. D.T.K., M.A., M.P., T.T.S.L developed nanosensors. K.V.V., B.J.R.S., I.K.Y.C., N.H.N., and H.J.C. generated plant materials and analyzed. X.G. performed image processing. T.K.P performed kinetics of fluorescent response. S.I.L., R.A., and G.P.S. generated materials for *in vitro* experiments. D.T.K., K.V.V., B.J.R.S., M.S.S., and I.C.J. wrote and edited the manuscript. All authors contributed to the analysis and approved the manuscript.

## References

1. Tim, W. & Joachim, von B. Climate Change Impacts on Global Food Security. Science (1979) 341, 508–513 (2013).

2. Xi, L., Zhang, M., Zhang, L., Lew, T. T. S. & Lam, Y. M. Novel Materials for Urban Farming. Advanced Materials 2105009 (2021) 10.1002/adma.202105009.

3. Teale, W. D., Paponov, I. A. & Palme, K. Auxin in action: signalling, transport and the control of plant growth and development. Nat Rev Mol Cell Biol 7, 847–859 (2006).

4. Bielach, A., Hrtyan, M. & Tognetti, V. B. Plants under stress: Involvement of auxin and cytokinin. Int J Mol Sci 18, (2017).

5. Friml, J. et al. Efflux-dependent auxin gradients establish the apical–basal axis of Arabidopsis. Nature 426, 147–153 (2003).

6. Benková, E. et al. Local, Efflux-Dependent Auxin Gradients as a Common Module for Plant Organ Formation. Cell 115, 591–602 (2003).

7. Blakeslee, J. J., Spatola Rossi, T. & Kriechbaumer, V. Auxin biosynthesis: spatial regulation and adaptation to stress. J Exp Bot 70, 5041–5049 (2019).

8. Enders, T. A. & Strader, L. C. Auxin activity: Past, present, and future. Am J Bot 102, 180–196 (2015).

9. Pařízková, B., Pernisová, M. & Novák, O. What Has Been Seen Cannot Be Unseen— Detecting Auxin In Vivo. Int J Mol Sci 18, (2017).

10. Abel, S. & Theologis, A. Early genes and auxin action. Plant Physiol 111, 9–17 (1996).

11. Kowalczyk, M. & Sandberg, G. Quantitative Analysis of Indole-3-Acetic Acid Metabolites in Arabidopsis. Plant Physiol 127, 1845–1853 (2001).

12. Tanaka, H., Dhonukshe, P., Brewer, P. B. & Friml, J. Spatiotemporal asymmetric auxin distribution: a means to coordinate plant development. Cell Mol Life Sci 63, 2738–2754 (2006).

13. Ulmasov, T., Murfett, J., Hagen, G. & Guilfoyle, T. J. Aux/IAA proteins repress expression of reporter genes containing natural and highly active synthetic auxin response elements. Plant Cell 9, 1963 (1997).

14. Ottenschläger, I. et al. Gravity-regulated differential auxin transport from columella to lateral root cap cells. Proceedings of the National Academy of Sciences 100, 2987–2991 (2003).

15. Brunoud, G. et al. A novel sensor to map auxin response and distribution at high spatio-temporal resolution. Nature 482, 103 (2012).

16. Herud-Sikimić, O. et al. A biosensor for the direct visualization of auxin. Nature 592, 768–772 (2021).

17. Okumoto, S., Jones, A. & Frommer, W. B. Quantitative Imaging with Fluorescent Biosensors. Annu Rev Plant Biol 63, 663–706 (2012).

18. Palmer, A. E., Qin, Y., Park, J. G. & McCombs, J. E. Design and application of genetically encoded biosensors. Trends Biotechnol 29, 144–152 (2011).

19. Blilou, I. et al. The PIN auxin efflux facilitator network controls growth and patterning in Arabidopsis roots. Nature 433, 39–44 (2005).

20. Matosevich, R. et al. Local auxin biosynthesis is required for root regeneration after wounding. Nat Plants 6, 1020–1030 (2020).

21. Qi, J. et al. Auxin depletion from leaf primordia contributes to organ patterning. Proceedings of the National Academy of Sciences 111, 18769–18774 (2014).

22. Deb, Y., Marti, D., Frenz, M., Kuhlemeier, C. & Reinhardt, D. Phyllotaxis involves auxin drainage through leaf primordia. Development 142, 1992–2001 (2015).

23. Koenig, D., Bayer, E., Kang, J., Kuhlemeier, C. & Sinha, N. Auxin patterns Solanum lycopersicum leaf morphogenesis. Development 136, 2997–3006 (2009).

24. Kodama, Y. Time Gating of Chloroplast Autofluorescence Allows Clearer Fluorescence Imaging In Planta. PLoS One 11, e0152484 (2016).

25. Donaldson, L. Autofluorescence in Plants. Molecules vol. 25 Preprint at 10.3390/molecules25102393 (2020).

26. Lew, T. T. S., Koman, V. B., Gordiichuk, P., Park, M. & Strano, M. S. The Emergence of Plant Nanobionics and Living Plants as Technology. Adv Mater Technol 5, 1900657 (2020).

27. Kwak, S.-Y. et al. Nanosensor Technology Applied to Living Plant Systems. doi:10.1146/annurev-anchem.

28. Lew, T. T. S. et al. Real-time detection of wound-induced H2O2 signalling waves in plants with optical nanosensors. Nat Plants 6, 404–415 (2020).

29. Lew, T. T. S., Park, M., Cui, J. & Strano, M. S. Plant Nanobionic Sensors for Arsenic Detection. Advanced Materials 33, 2005683 (2021).

30. Zhang, J. et al. Molecular recognition using corona phase complexes made of synthetic polymers adsorbed on carbon nanotubes. Nat Nanotechnol 8, 959 (2013).

31. Roig-Villanova, I. & Martínez-García, J. F. Plant responses to vegetation proximity: A whole life avoiding shade. Front Plant Sci 7, 1–10 (2016).

32. Boghossian, A. A. et al. Near-Infrared Fluorescent Sensors based on Single-Walled Carbon Nanotubes for Life Sciences Applications. ChemSusChem 4, 848–863 (2011).

33. Tan, X. et al. Mechanism of auxin perception by the TIR1 ubiquitin ligase. Nature 446, 640 (2007).

34. Dharmasiri, N., Dharmasiri, S. & Estelle, M. The F-box protein TIR1 is an auxin receptor. Nature 435, 441–445 (2005).

35. Kepinski, S. & Leyser, O. The Arabidopsis F-box protein TIR1 is an auxin receptor. Nature 435, 446–451 (2005).

36. Park, J. et al. Molecular interactions of polyimides with single-walled carbon nanotubes. Polym Chem 4, 290–295 (2013).

37. Ljung, K., Bhalerao, R. P. & Sandberg, G. Sites and homeostatic control of auxin biosynthesis in Arabidopsis during vegetative growth. The Plant Journal 28, 465–474 (2001).

38. Giraldo, J. P. et al. A Ratiometric Sensor Using Single Chirality Near-Infrared Fluorescent Carbon Nanotubes: Application to in Vivo Monitoring. Small 11, 3973–3984 (2015).

39. Leasure, C. D., Chen, Y. P. & He, Z. H. Enhancement of Indole-3-Acetic acid photodegradation by vitamin B6. Molecular Plant vol. 6 1992–1995 Preprint at 10.1093/mp/sst089 (2013).

40. Skalický, V. et al. Auxin metabolite profiling in isolated and intact plant nuclei. Int J Mol Sci 22, (2021).

41. Skalický, V., Kubeš, M., Napier, R. & Novák, O. Auxins and cytokinins—The role of subcellular organization on homeostasis. International Journal of Molecular Sciences vol. 19 Preprint at 10.3390/ijms19103115 (2018).

42. Geisler, M. M. A Retro-Perspective on Auxin Transport. Front Plant Sci 12, (2021).

43. Wong, M. H. et al. Lipid Exchange Envelope Penetration (LEEP) of Nanoparticles for Plant Engineering: A Universal Localization Mechanism. Nano Lett 16, 1161–1172 (2016).

44. Zuo, J., Niu, Q.-W. & Chua, N.-H. An estrogen receptor-based transactivator XVE mediates highly inducible gene expression in transgenic plants. The Plant Journal 24, 265–273 (2000).

45. Vanneste, S. & Friml, J. Auxin: A Trigger for Change in Plant Development. Cell 136, 1005–1016 (2009).

46. Olivier, M., Anne-Sophie, F., Ioannis, X. & Christian, F. Local auxin production underlies a spatially restricted neighbor-detection response in Arabidopsis. Proceedings of the National Academy of Sciences 114, 7444–7449 (2017).

47. Adamowski, M. & Friml, J. PIN-Dependent Auxin Transport: Action, Regulation, and Evolution. Plant Cell 27, 20–32 (2015).

48. Lew, T. T. S. et al. Species-independent analytical tools for next-generation agriculture. Nat Plants 6, 1408–1417 (2020).

49. Nakamura, A. et al. Brassinolide Induces IAA5, IAA19 and DR5, a Synthetic Auxin Response Element in Arabidopsis, Implying a Cross Talk Point of Brassinosteroid and Auxin Signaling. Plant Physiol. 133, 1843–1853 (2003).

50. Nemhauser, J. L., Mockler, T. C. & Chory, J. Interdependency of Brassinosteroid and Auxin Signaling in Arabidopsis. PLOS Biol. 2, e258 (2004).

51. Korasick, D. A., Enders, T. A. & Strader, L. C. Auxin biosynthesis and storage forms. J. Exp. Bot. 64, 2541–2555 (2013).

52. Yang, J. et al. Dynamic Regulation of Auxin Response during Rice Development Revealed by Newly Established Hormone Biosensor Markers. Frontiers in Plant Science vol. 8 Preprint at (2017).

53. Chen, Y., Yordanov, Y. S., Ma, C., Strauss, S. & Busov, V. B. DR5 as a reporter system to study auxin response in Populus. Plant Cell Rep. 32, 453–463 (2013).

54. Almeida Trapp, M., De Souza, G. D., Rodrigues-Filho, E., Boland, W. & Mithöfer, A. Validated method for phytohormone quantification in plants. Front. Plant Sci. vol. 5 417 Preprint at (2014).

55. Matsuda, F., Miyazawa, H., Wakasa, K. & Miyagawa, H. Quantification of Indole-3-Acetic Acid and Amino Acid Conjugates in Rice by Liquid Chromatography– Electrospray Ionization–Tandem Mass Spectrometry. Biosci Biotechnol Biochem 69, 778–783 (2005).

56. Xiong, Y. & Jiao, Y. The diverse roles of auxin in regulating leaf development. Plants vol. 8 Preprint at 10.3390/plants8070243 (2019).

57. Chun-Yi Ang, M., et al. Nanosensor Detection of Synthetic Auxins In Planta using Corona Phase Molecular Recognition. ACS Sens 6, 3032–3046 (2021).

58. Jorio, A. & Saito, R. Raman spectroscopy for carbon nanotube applications. J Appl Phys 129, (2021).

59. Cao, Y. et al. Drug Delivery in Plants Using Silk Microneedles. Advanced Materials 35, (2023).

60. Kwak, S.-Y. et al. A Nanobionic Light-Emitting Plant. Nano Lett 17, 7951–7961 (2017).

61. Boonyaves, K. et al. Near-Infrared Fluorescent Carbon Nanotube Sensors for the Plant Hormone Family Gibberellins. Nano Lett 23, 916–924 (2023).

62. Sng, B. J. R. et al. Rapid metabolite response in leaf blade and petiole as a marker for shade avoidance syndrome. Plant Methods 16, (2020).

63. Creely, C. M., Singh, G. P. & Petrov, D. Dual wavelength optical tweezers for confocal Raman spectroscopy. Opt Commun 245, 465–470 (2005).

