## Supplementary material for "A Near Infrared Fluorescent Nanosensor for Spatial and Dynamic Measurements of Auxin, Indole-3-Acetic Acid, *in Planta*"

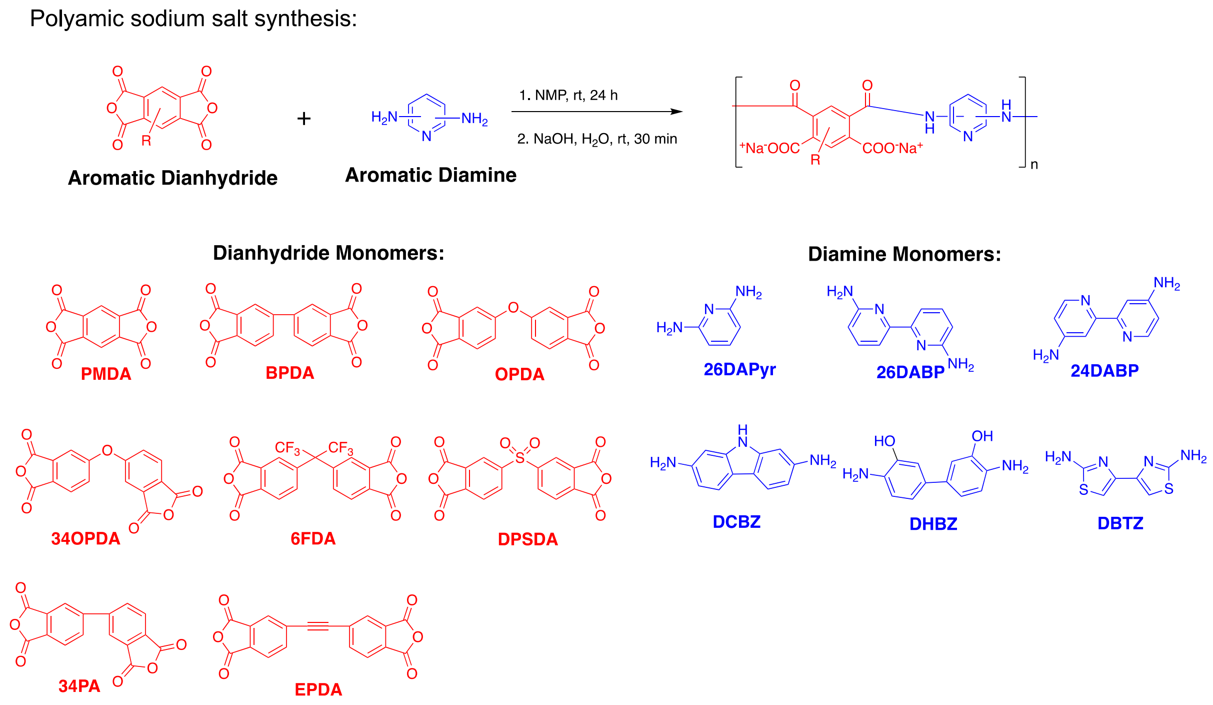

**Supplementary Fig. 1. Synthesis of polyamic sodium salts.** Synthesis scheme of a series polyamic acid sodium salt analogues by combination with different dianhydride and diamine monomers. NMP: N-methyl-2-pyrrolidone; rt: room temperature; PMDA, pyromellitic dianhydride; BPDA, 3,3',4,4'-biphenyltetracarboxylic dianhydride; OPDA, 4,4'-oxydiphthalic anhydride; 34OPDA, 3,4'-oxydiphthalic anhydride; 6FDA, 4,4'-(hexafluoroisopropylidene)diphthalic anhydride, DPSDA, 3,3',4,4'-diphenylsulfonetetracarboxylic dianhydride; 34PA, 3,4'-bithphalic anhydride; EPDA, 4,4'-(ethyne-1,2-diyl)diphthalic anhydride; 26DAPyr, 2,6-diaminopyridine; 26DABP, 6,6'-diamino-2,2'-bipyridyl; 24DABP, 4,4'-diamino-2,2'-bipyridyl; DCBZ, 3,6-diaminocarbazole; DHBZ, 3,3'-dihydroxybenzidine; DBTZ, 2,2-diamino-4,4'-bithiazole.

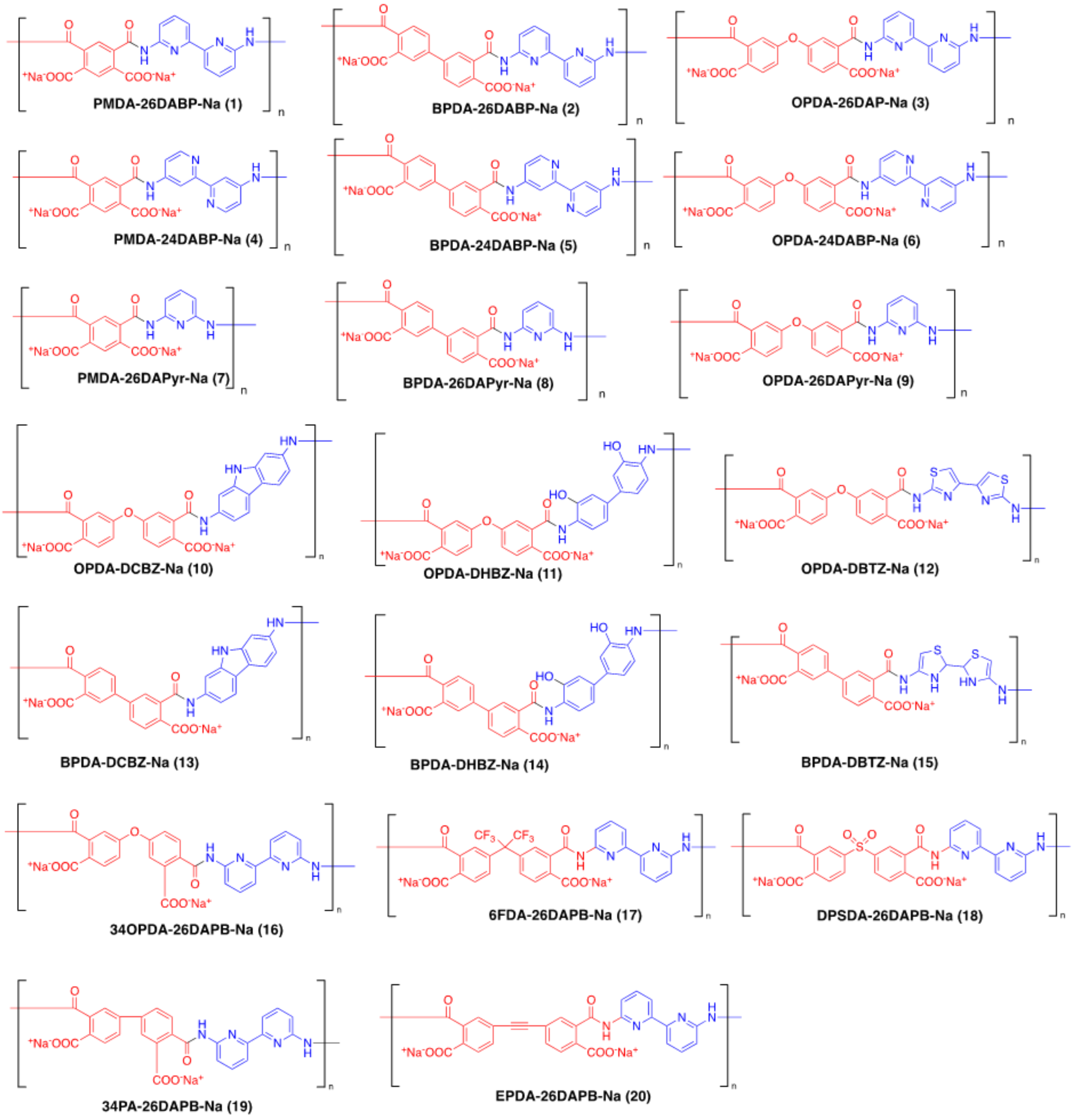

**Supplementary Fig. 2. Polymer structures of polyamic acid sodium salts used in the suspension of SWNTs for IAA screening.** Abbreviations are listed in Supplementary Fig. 1.

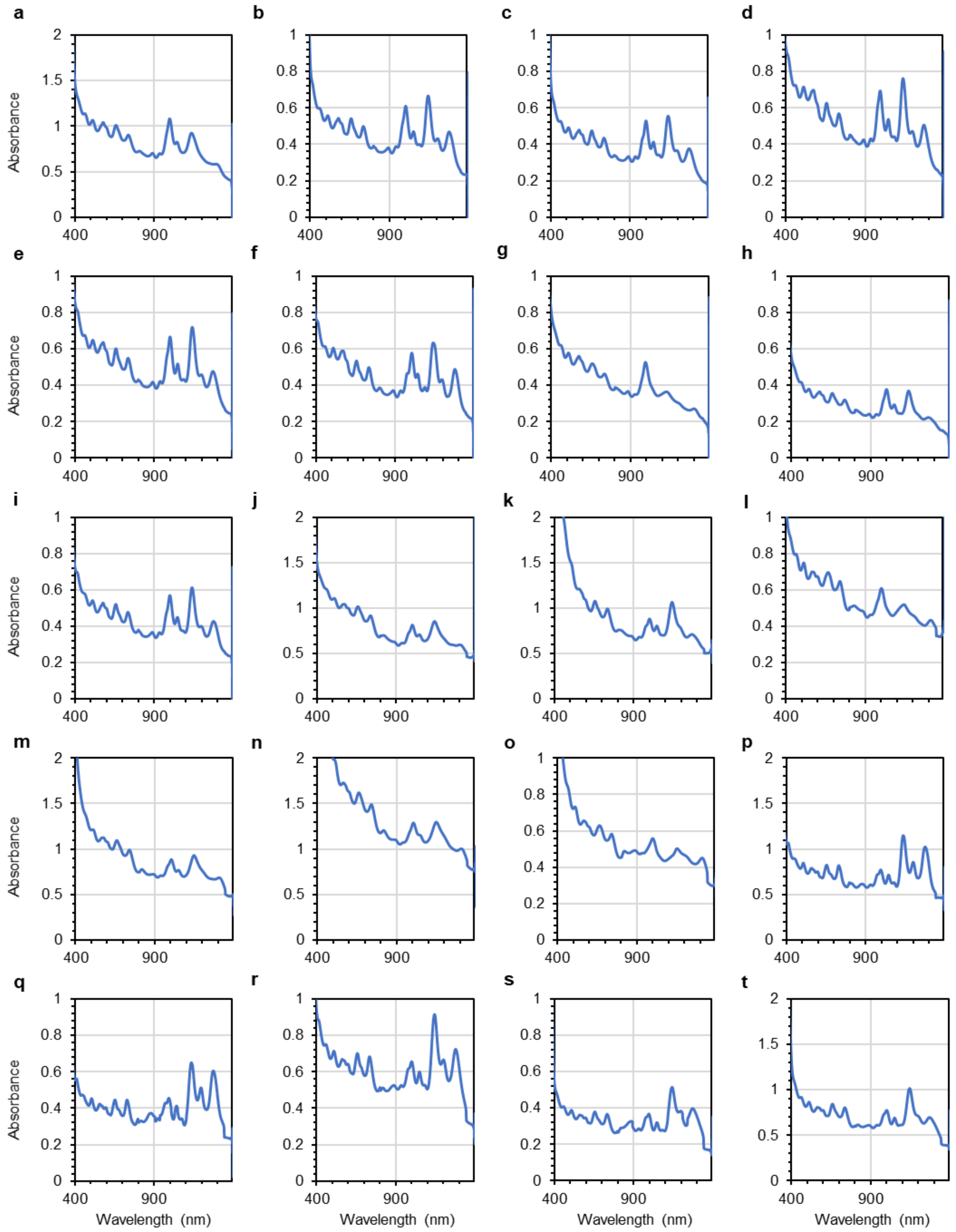

**Supplementary Fig. 3. UV-vis-nIR absorption spectrum of CoPhMoRe candidates for IAA screening.** **(a)** PMDA-26DABP-Na-SWNT. **(b)** PMDA-24DABP-Na-SWNT. **(c)** PMDA-26DAPyr-Na-SWNT. **(d)** BPDA-26DABP-Na-SWNT. **(e)** BPDA-24DABP-Na-SWNT. **(f)** BPDA-26DAPyr-Na-SWNT. **(g)** OPDA-26DABP-Na-SWNT. **(h)** OPDA-24DABP-Na-SWNT. **(i)** OPDA-26DAPyr-Na-SWNT**. (j)** OPDA-DCBA-Na-SWNT. **(k)** OPDA-DHBZ-Na-SWNT. **(l)** OPDA-DBTZ-Na-SWNT. **(m)** BPDA-DCBA-Na-SWNT. **(n)** BPDA-DHBZ-Na-SWNT. **(o)** BPDA-DBTZ-Na-SWNT. **(p)** 34OPDA-26DABP-Na-SWNT. **(q)** 6FDA-26DABP-Na-SWNT. **(r)** DPSDA-26DABP-Na-SWNT. **(s)** 34PA-26DABP-Na-SWNT. **(t)** EPDA-26DABP-Na-SWNT. Abbreviations are listed in Supplementary Fig. 1.

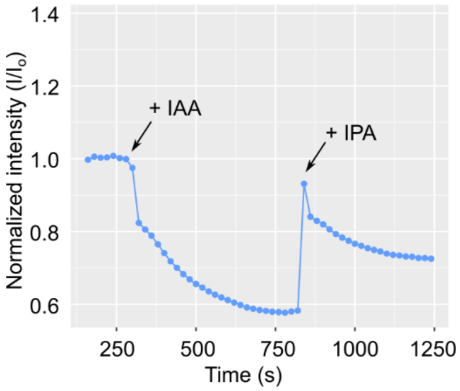

**Supplementary Fig. 4. Binding competition between IAA and IPA.** Kinetic response of IAA nanosensor in the presence of both IAA and IPA at 1:10 molar ratio. Initial addition of IAA to the IAA nanosensor resulted in fluorescence quenching. Subsequent adding of IPA resulted in an immediate turn-on response. The turn-on signal then subsequently quenched to the new equilibrium state, which depends on the ratio between IAA and IPA in the solution. This observation indicates that IPA has fast binding kinetics to the IAA nanosensor, but binding of IAA to the nanosensor is more thermodynamically favorable. Three independent meaurements were repeated with similar results (*n*=3). IAA, indole-3-acetic acid; IPA, indole-3-pyruvic acid. Arrows indicate time-points of adding IAA and IPA.

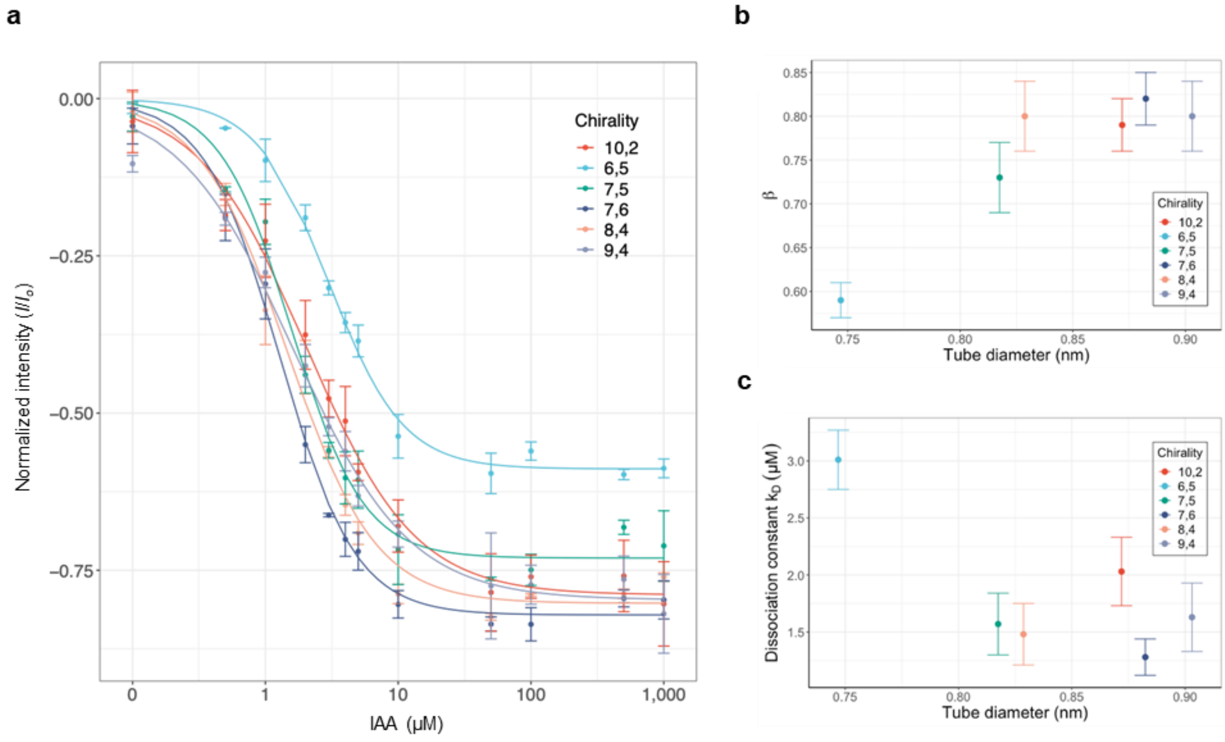

**Supplementary Fig. 5. Effect of SWNT chirality on IAA sensing. (a)** Calibration curves of sensor response versus IAA concentration for SWNT chiralities (6,5), (8,4), (7,5), (7,6), (9,4), and (10,2) and their corresponding fits. **(b, c)** Scatter plots representing two fit parameters β and $K_d$ from Langmuir isotherm model versus different SWNT chiralities in terms of tube diameters. In **a**−**c**, the error bars are standard deviation obtained from three different measurements (*n*=3).

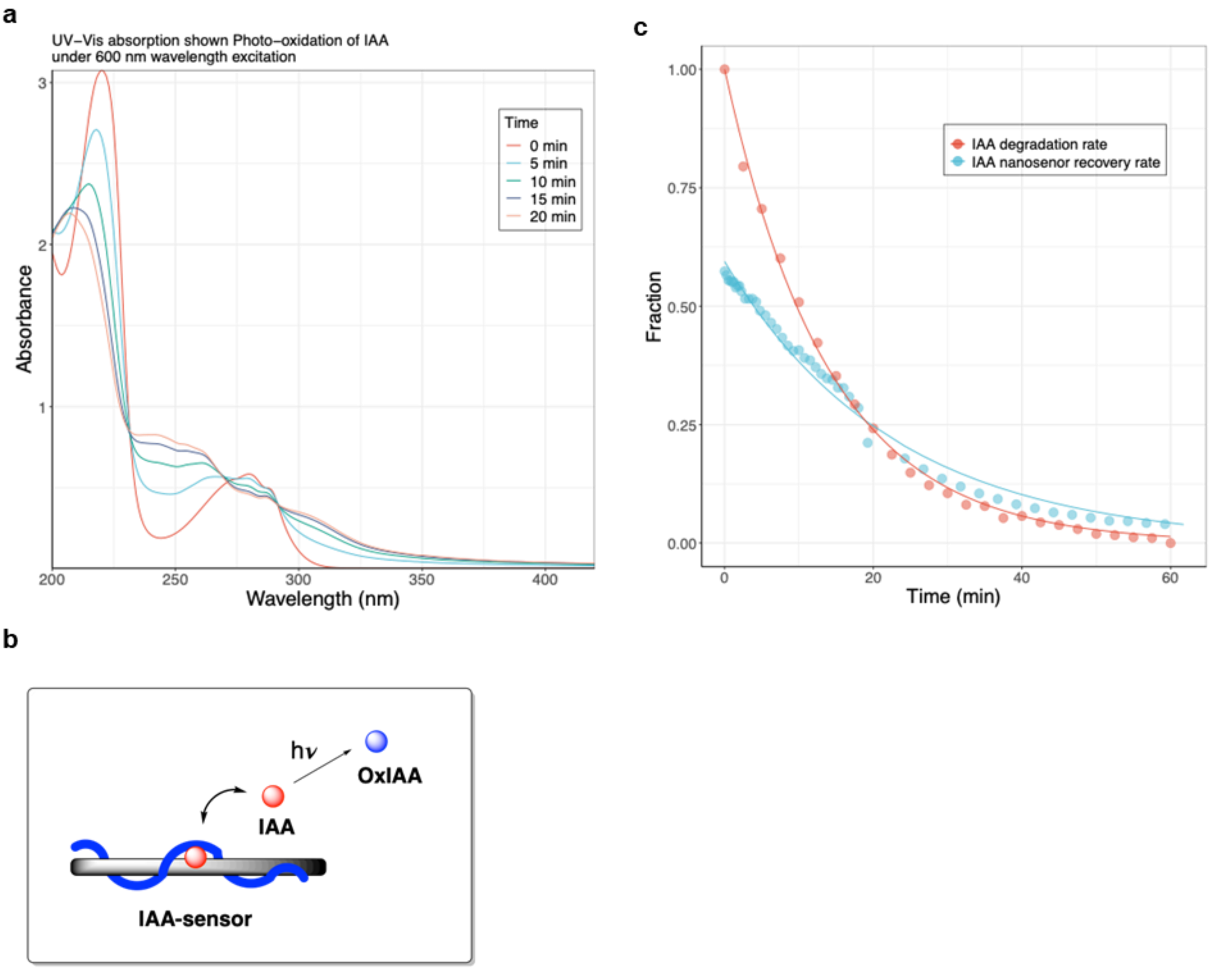

**Supplementary Fig. 6. Sensor responsiveness based on in situ IAA photodegradation study. (a)** Degradation of IAA due to prolonged exposure to 600 nm excitation source as showed by UV-Vis absorption. **(b)** Simplified model showed the binding activity of IAA nanosensor to IAA during photodegradation. **(c)** Comparison between kinetic response of IAA sensor to IAA (as shown by fluorescent recovery kinetics, cyan line) and IAA degradation rate (red line). IAA, indole-3-acetic acid; OxIAA, 2-oxindole-3-acetic acid.

### Kinetics of fluorescent response to auxin photo-degradation

To study the kinetics of fluorescent response to auxin photo-degradation, a mathematical model was developed assuming the relationship between the corona phase binding sites and auxin under photodegradation based on the diagram in Figure S7a:

$$\theta - IAA \leftrightarrow \theta + IAA \overset{h\nu}{\rightarrow} OxIAA \quad (1)$$

Where the equation rates for concentration occupied binding sites $[\theta - IAA]$ and IAA can be written as:

$$\frac{d[\theta - IAA]}{dt} = k_a[\theta][IAA] - k_d[\theta - IAA] \quad (2)$$

$$\frac{d[IAA]}{dt} = -k_a[\theta][IAA] + k_d[\theta - IAA] - k_{ox}[IAA] \quad (3)$$

Where $k_a$ and $k_d$ are the rate constants of association and dissociation respectively of binding process and $k_{ox}$ is the degradation rate constant of oxidation of IAA which is an irreversible process. Assuming a pseudo-equilibrium condition for free IAA where the extent of IAA dissociated from corona phase is simultaneously degraded by photo-oxidation such that $\frac{d[IAA]}{dt} = 0$, free IAA concentration can be estimated as:

$$[IAA] = \frac{k_d[\theta - IAA]}{k_a[\theta] + k_{ox}} \quad (4)$$

By substituting (4) into (2):

$$\frac{d[\theta - IAA]}{dt} = k_a[\theta]\frac{k_d[\theta - IAA]}{k_a[\theta] + k_{ox}} - k_d[\theta - IAA]$$

$$\frac{d[\theta - IAA]}{dt} = k_d[\theta - IAA]\left(\frac{k_a[\theta]}{k_a[\theta] + k_{ox}} - 1\right)$$

Since $[\theta]_{tot} = [\theta - IAA] + [\theta]$,

$$\frac{d[\theta - IAA]}{dt} = k_d[\theta - IAA]\left(\frac{k_a([\theta]_{tot} - [\theta - IAA])}{k_a([\theta]_{tot} - [\theta - IAA]) + k_{ox}} - 1\right)$$

$$\frac{d[\theta - IAA]}{dt} = k_d[\theta - IAA]\left(\frac{k_a([\theta]_{tot} - [\theta - IAA]) - k_a([\theta]_{tot} - [\theta - IAA]) - k_{ox}}{k_a([\theta]_{tot} - [\theta - IAA]) + k_{ox}}\right)$$

$$\frac{d[\theta - IAA]}{dt} = k_d[\theta - IAA]\left(\frac{k_a[\theta]_{tot} - k_a[\theta - IAA] - k_a[\theta]_{tot} + k_a[\theta - IAA] - k_{ox}}{k_a([\theta]_{tot} - [\theta - IAA]) + k_{ox}}\right)$$

$$\frac{d[\theta - IAA]}{dt} = \frac{-k_d k_{ox}[\theta - IAA]}{(k_a[\theta]_{tot} + k_{ox}) - k_a[\theta - IAA]}$$

$$dt = \frac{-(k_a[\theta]_{tot} + k_{ox}) + k_a[\theta - IAA]}{k_d k_{ox}[\theta - IAA]} d[\theta - IAA]$$

Assuming that at $t = 0, [\theta - IAA] = [\theta - IAA]_o$, integrating both sides gives:

$$\int_0^t dt = \frac{1}{k_d k_{ox}} \int_{[\theta-IAA]_0}^{[\theta-IAA]} \left(k_a - \frac{(k_a[\theta]_{tot} + k_{ox})}{[\theta - IAA]}\right) d[\theta - IAA]$$

$$t = \frac{1}{k_d k_{ox}} \left[k_a([\theta - IAA] - [\theta - IAA]_0) - (k_a[\theta]_{tot} + k_{ox}) ln \frac{[\theta - IAA]}{[\theta - IAA]_0}\right]$$

At early time when t = to, $[\theta - IAA] - [\theta - IAA]_0 \approx 0$

At time increases when $[\theta - IAA] \ll [\theta - IAA]_o$,we have $ln \frac{[\theta-IAA]}{[\theta-IAA]_0} \gg [\theta - IAA] - [\theta - IAA]_0$ and $(k_a[\theta]_{tot} + k_{ox}) > k_a$, we can approximate that

$$k_a([\theta - IAA] - [\theta - IAA]_0) - (k_a[\theta]_{tot} + k_{ox}) ln \frac{[\theta-IAA]}{[\theta-IAA]_0} \approx -(k_a[\theta]_{tot} + k_{ox}) ln \frac{[\theta-IAA]}{[\theta-IAA]_0}$$

Therefore, $t = -\frac{k_a[\theta]_{tot}+k_{ox}}{k_d k_{ox}} ln \frac{[\theta-IAA]}{[\theta-IAA]_0}$

or

$$\frac{[\theta - IAA]}{[\theta - IAA]_0} = e^{-\frac{k_d k_{ox}}{k_a[\theta]_{tot}+k_{ox}} t} = e^{-k' t}$$

Therefore, we can obtain the time-dependence concentration of $[\theta - IAA]$

$$\frac{I - I_o}{I_o} = -\beta \frac{[\theta - IAA]}{[\theta - IAA]_0} \cong -\beta e^{-\frac{k_d k_{ox}}{k_a[\theta]_{tot}+k_{ox}} t} = -\beta e^{-k' t} \quad (5)$$

Where β is proportionality factor. By fitting the model to the time-dependent fluorescent response data, β and apparent constant $k'$ were determined as 0.59 ± 0.1 and 0.043 ± 0.02 ($min^{-1}$) respectively. The proportionality factor β obtained here is similar to the proportionality factor β obtained from the calibration curve of SWNT (6,5). This is expected because at 600 nm excitation wavelength, fluorescent of (6,5) is the brightest.

From $k' = \frac{k_d}{k_a[\theta]_{tot}+k_{ox}} k_{ox}$, it can be implied that the responsiveness of auxin sensor **(3)** could be improved such that $k' \approx k_{ox}$when the dissociation constant $k_d \approx k_a[\theta]_{tot} + k_{ox}$. While $k_{ox}$ is an intrinsic property of IAA, the dissociation/ association constants, $k_d$ and $k_{a,}$ can be controlled by using different SWNT chiralities and the concentration of total binding sites $[\theta]_{tot}$ is a variable that highly dependent on the surface coverage of adsorbed polymer on nanoparticle surface. Therefore, this kinetic model proposes an opportunity for improvement of sensor sensitivity and responsiveness by tuning the concentration of binding sites as well as appropriate choice of SWNT chirality.

Likewise, the kinetic of photodegradation of IAA can be simplified by the following irreversible reaction: $IAA \overset{h\nu}{\rightarrow} oxIAA$; assuming that this is a first order kinetic process, the time dependence concentration of IAA can be written as:

$$ln\left(\frac{[IAA]_t}{[IAA]_{t=0}}\right) = -k_{ox}t \tag{1}$$

Where $k_{ox}$ is degradation rate constant. Because the nIR fluorescent intensity almost returned to its initial intensity after one hour, we may assume that IAA is almost completely degraded ($[IAA]_{t=60} \cong 0$) at t = 60 min. Assuming that the IAA concentration is proportional to its absorbance value $Abs_t$ at 280 nm, we can derive that:

$$ln\left(\frac{Abs_t - Abs_{t=60}}{Abs_{t=0} - Abs_{t=60}}\right) = ln\left(\frac{[A]}{[A]_o}\right) = -k_{ox}t \tag{2}$$

Where $Abs_{t=60}$ accounts for absorbance value background when [IAA] = 0. Value of $k_{ox}$ was obtained as 0.071 min$^{-1}$ by fitting to time-dependence absorbance value at 280 nm, where orange line is the fitting curve to blue data points (calculated from absorbance values as shown in equation (2).

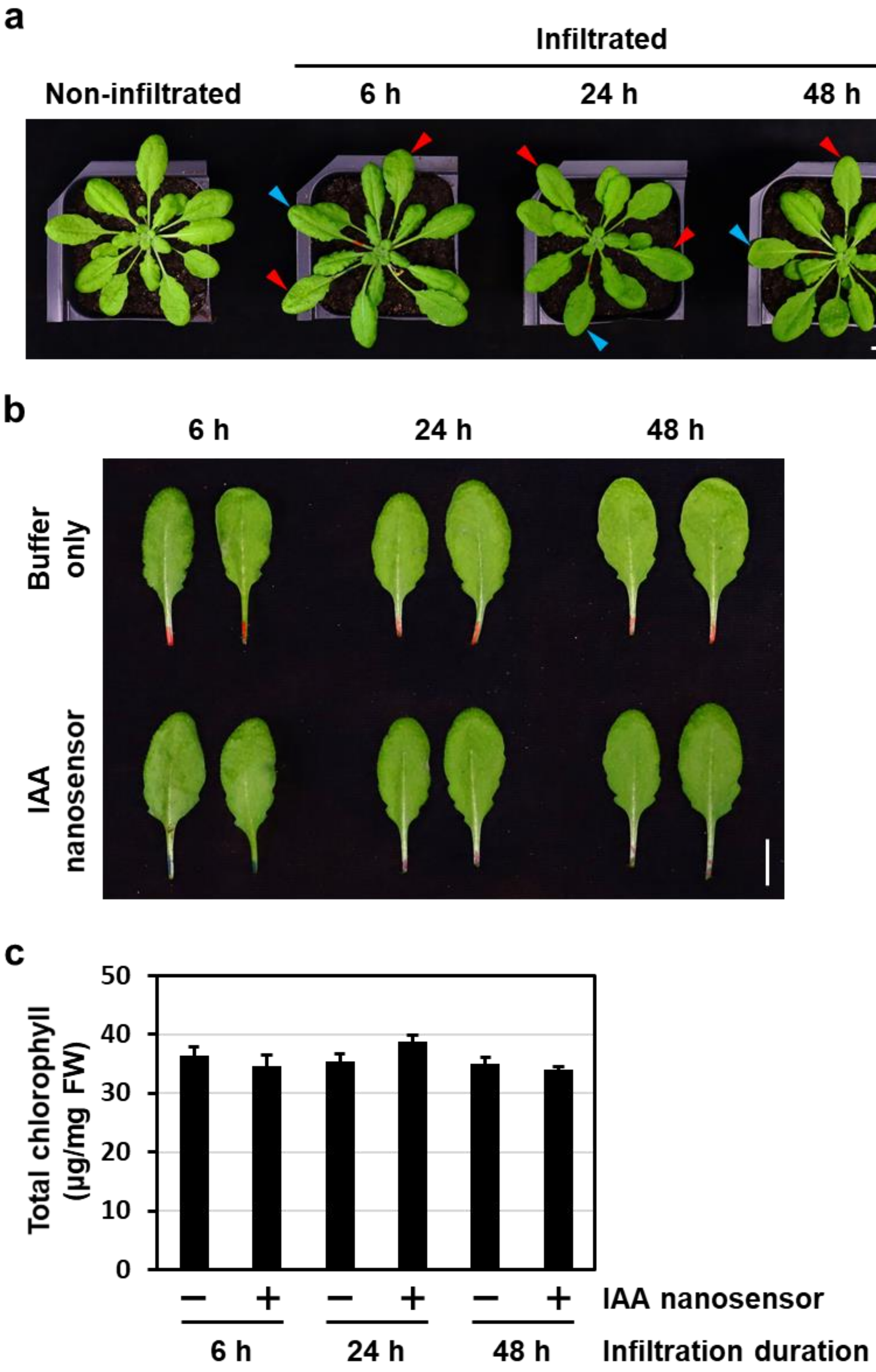

**Supplementary Fig. 7. Phenotype and chlorophyll content of Arabidopsis leaves infiltrated with or without IAA nanosensor. (a)** Phenotype of Arabidopsis plants without or with infiltration. Infiltrated plants were observed at 6, 24, and 48 h post-infiltration. Red arrowhead, leaf infiltrated with IAA nanosensor. Blue arrowhead, leaf infiltrated with 10 mM MES buffer (pH 5.5). Scale, 3 cm. **(b)** Dissected leaves infiltrated with IAA nanosensor or MES buffer in **(a)**. Scale, 1 cm. **(c)** Total chlorophyll content of leaves infiltrated with IAA nanosensor or MES buffer in **(b)**. *n*=3. Values represent mean ± standard deviation.

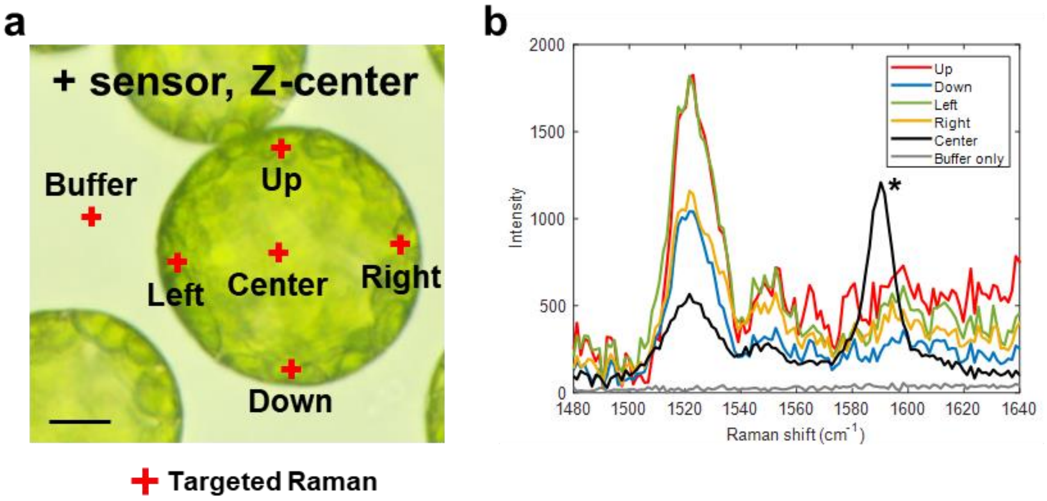

**Supplementary Fig. 8. IAA nanosensor localized towards the center of Arabidopsis protoplast. (a)** Bright field image of protoplast isolated from Arabidopsis leaf blade infiltrated with the IAA nanosensor. Optical-sectioning focal plane was set at the center of the protoplast (Z-center). Locations indicated by a red cross were subjected to confocal Raman spectroscopy. Scale, 10 µm. **(b)** Raman spectra of protoplast locations in **a**, showing 1525 $cm^{-1}$ carotenoid peak and 1590 $cm^{-1}$ IAA nanosensor G-band.

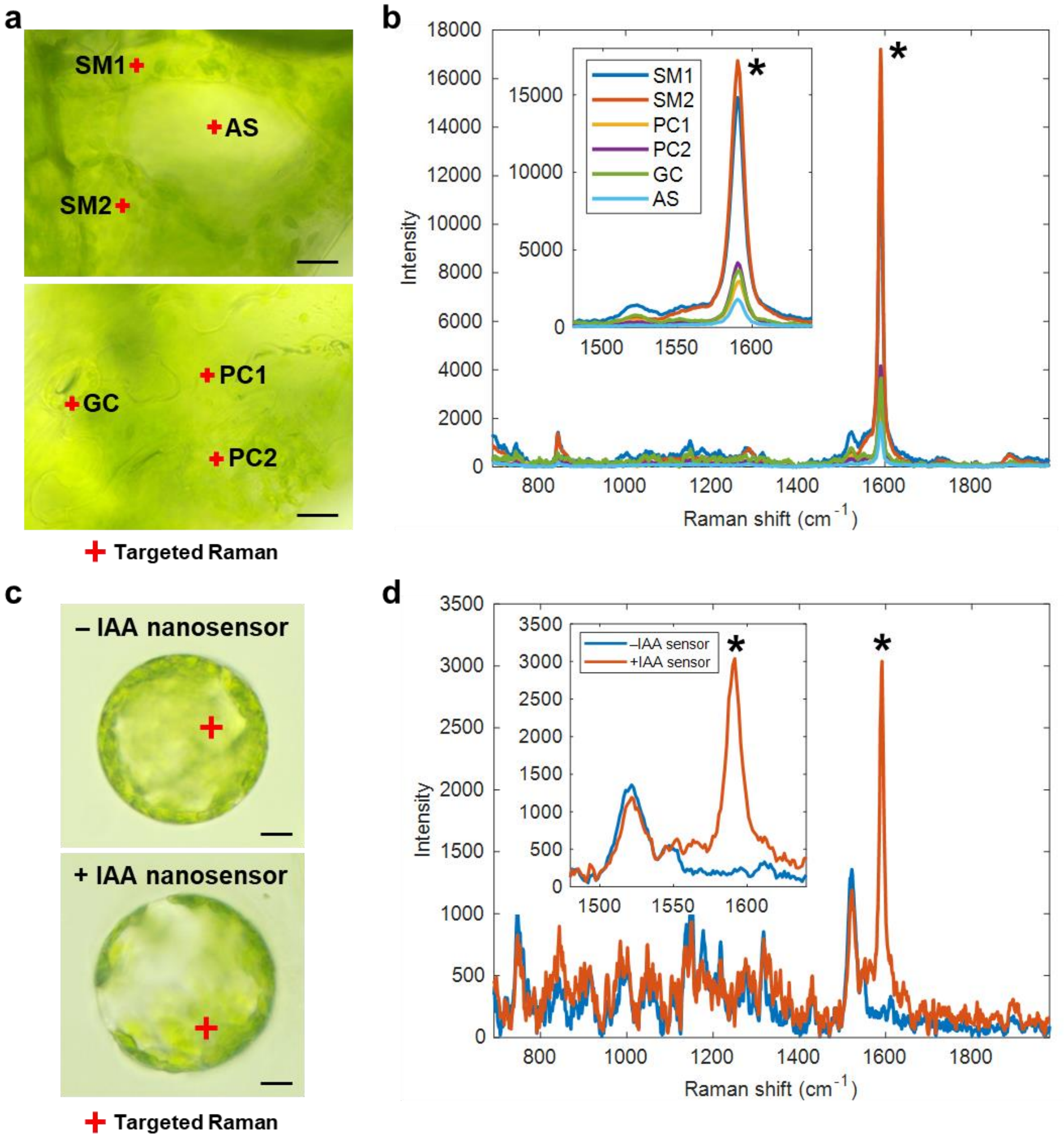

**Supplementary Fig. 9. IAA nanosensor localizes into *Nicotiana benthamiana* leaf cells after infiltration. (a)** Bright field images of cells in *N. benthamiana* leaf blade after infiltration with IAA nanosensor. Top, transverse cross-section. Bottom, abaxial leaf epidermis. Locations indicated by a red cross were subjected to confocal Raman spectroscopy. Scale, 20 µm. **(b)** Raman spectra of individual leaf cells and structure in **a**. Inset, zoomed in Raman spectra showing 1590 $cm^{-1}$ IAA nanosensor G-band. **(c)** Protoplasts isolated from *N. benthamiana* with and without IAA nanosensor infiltration. Locations indicated by a red cross were subjected to confocal Raman spectroscopy. Scale, 10 µm. **(d)** Average Raman spectra of protoplasts in **c**. $n = 4$. Inset, zoomed in Raman spectra showing 1525 $cm^{-1}$ carotenoid peak and 1590 $cm^{-1}$ IAA nanosensor G-band. Asterisk (*) indicates IAA nanosensor G-band at 1590 $cm^{-1}$ Raman shift.

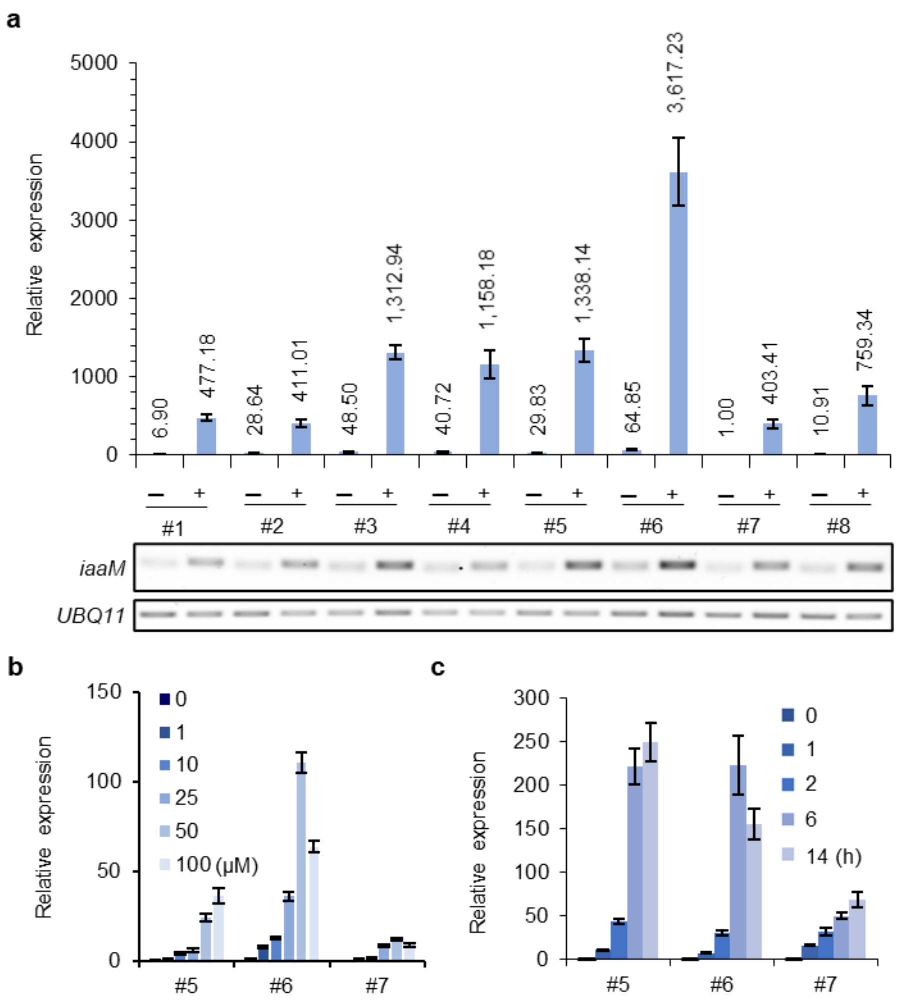

**Supplementary Fig. 10. Inducible expression of iaaM by varying β-estradiol concentrations and time course induction in XVE::iaaM Arabidopsis plants. (a)** 10-d-old seedlings of different *XVE::iaaM* Arabidopsis transgenic T2 lines were subjected to 50 μM β-estradiol induction (+) overnight (16 h) under continuous light. Expression of *iaaM* at different concentrations of β-estradiol (0−100 μM) for 6 h. **(b)** or at different time points with 50 μM β-estradiol. (**c**) under continuous light in *XVE::iaaM* transgenic Arabidopsis lines (#5, 6, and 7). Data are mean ± s.d. from three biological replicates (*n*=3).

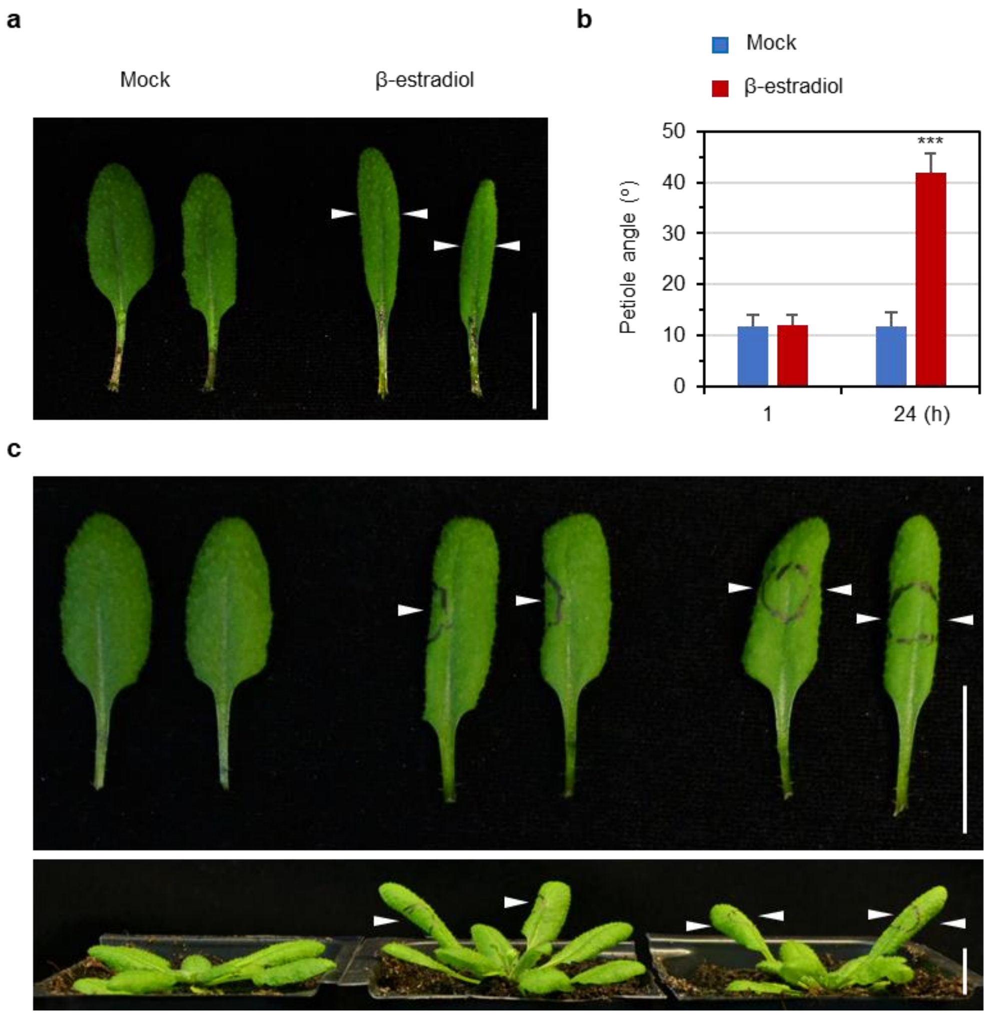

**Supplementary Fig. 11. Auxin-induced phenotypic changes of *XVE::iaaM* Arabidopsis leaves under β-estradiol treatment.** 4-w-old Arabidopsis plants expressing *XVE::iaaM* were applied with DMSO (mock) or 100 μM β-estradiol using a paintbrush to induce the expression of *iaaM*. **(a)** Rosette leaves after 24 h of mock or β-estradiol treatment on the whole leaf blade. Arrow heads represent curled leaves. Scale bar, 1 cm. **(b)** Measurement of leaf hyponasty after 24 h of β-estradiol treatment. Data are mean ± s.d. (*n*=15). ***p < 0.001. **(c)** Local β-estradiol. Top panel indicates detached leaves while bottom panel indicates the whole plants. Left (no treatment), middle (side treatment), and right (middle leaf treatment). Black circles indicate positions of local β-estradiol treatment. Arrow heads represent curled leaves. 10 independent biological experiments were repeated with similar results (*n*=5). Scale bars, 1 cm**.**

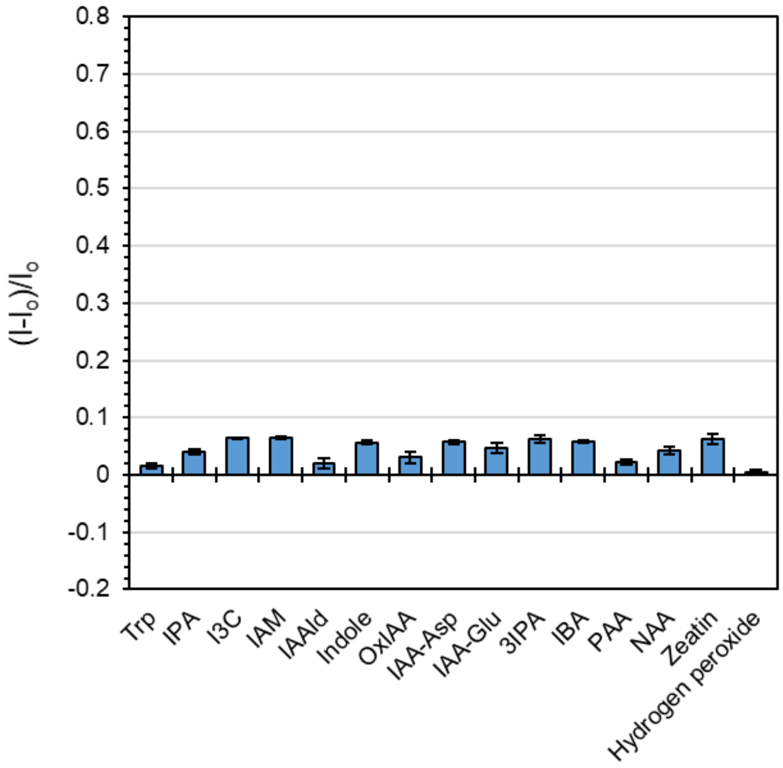

**Supplementary Fig. 12. Optical modulation of reference sensor $(AT)_{15}$-SWNT against plant analytes.** The screening was carried out using 1 mg/L (AT)15-SWNT in 10 mM MES buffer pH 5.5 with 100 µM of respective plant analytes. The error bars represent standard deviation of triplicate.

**Supplementary Table 1** Primers used in the study for *iaaM* cloning and qRT-PCR

| Primer name | Sequence (5’ to 3’) | Function |
|---|---|---|
| iaaM-F | AAAAAGCAGGCTTCATGTATGATCATTTCAACTC | *iaaM* cloning with attb adapters for Gateway cloning |
| iaaM-R | AGAAAGCTGGGTGTCAGTATCTATAAGAAGCGTTGA | |
| attb1-F | GGGGACAAGTTTGTACAAAAAAGCAGGCT | |
| attb2-R | GGGGACCACTTTGTACAAGAAAGCTGGGT | |
| iaaM qPCR-F | GATTGGAAGAGGCTGCTATCG | *iaaM* expression analysis |
| iaaM qPCR-R | GTCTAGCCCATCTATCTCCTCC | |
| GH3.3 qPCR-F | ACAATTCCGCTCCACAGTTC | *GH3.3* expression analysis |
| GH3.3 qPCR-R | ACGAGTTCCTTGCTCTCCAA | |
| NbGH3.6 qPCR-F | GCTATTAGCAATGGCGCTTC | *NbGH3.6* expression analysis |
| NbGH3.6 qPCR-R | ATTGCTTGTGACCAGGAACC | |
| Actin2 qPCR-F | GATCTCCAAGGCCGAGTATGAT | Internal control for expression analysis |
| Actin2 qPCR-R | CCCATTCATAAAACCCCAGC | |

**Reversible first-order binding model for IAA nanosensor.**
The binding event between IAA nanosensor (S) and IAA may be assumed as a first-order reversible reaction:

$$IAA + S \underset{k_b}{\overset{k_f}{\rightleftharpoons}} S - IAA$$

- $k_f$ and $k_b$ are forward and backward rate constants.

The rate of reaction can be described by the following equation:

$$\frac{d[S]}{dt} = -k_f[S][IAA] + k_b[S - IAA]$$

At the initial stage of binding events, the sensor and the analyte are close to their original concentrations, $[S] \approx [S]_o$ ,$[IAA] \approx [IAA]_o$ and $[S - IAA] \approx 0$, therefore:

$$\left.\frac{d[S]}{dt}\right|_{t\to 0} = -k_f[S]_0[IAA]_o$$

Assuming that $I/I_o = [S]/[S]_o$, therefore:

$$\frac{d[S]}{dt} = [S]_o\frac{d([S]/[S]_o)}{dt} = [S]_o\frac{d(I/I_o)}{dt}$$

$$\left.\frac{d(I/I_o)}{dt}\right|_{t\to 0} [S]_o = -k_f[S]_0[IAA]_o$$

$$\left.\frac{d(I/I_o)}{dt}\right|_{t\to 0} = -k_f[IAA]_o$$

Therefore, the forward reaction rate is

$$k_f = \frac{-1}{[IAA]_0} \times \left.\frac{d(I/I_o)}{dt}\right|_{t\to 0} \qquad (1)$$

At equilibrium, we have

$$\frac{d[S]}{dt} = -k_f[S][IAA] + k_b[S - IAA] = 0$$

Therefore,

$$k_b = \frac{k_f[S][IAA]}{[S - IAA]}$$

From mass balance equation:

$$[S]_0 = [S] + [S - IAA] = constant$$

Therefore,

$$\frac{I}{I_0} = \frac{[S]}{[S]_0} = \frac{[S]_0 - [S - IAA]}{[S]_0} = 1 - \frac{[S - IAA]}{[S]_0}$$

$$[S-IAA]=\left(1-\frac{I}{I_0}\right)[S]_0$$

Therefore,

$$k_b=\frac{k_f[IAA]\left({}^{I}/{}_{I_o}\right)}{(1-{}^{I}/{}_{I_0})} \quad (2)$$

The concentration [IAA] can then be approximated

$$\frac{d[S]}{dt}=-k_f[S][IAA]+k_b[S-IAA]$$

$$[IAA]=\frac{\frac{d[S]}{dt}-k_b[S-IAA]}{-k_f[S]}$$

$$[IAA]=\frac{[S]_o\frac{d({}^{I}/{}_{I_o})}{dt}-k_b\left(1-\frac{I}{I_0}\right)[S]_0}{-k_f\frac{I}{I_0}[S]_0}$$

$$[IAA]=\frac{k_b\left(1-\frac{I}{I_0}\right)-\frac{d({}^{I}/{}_{I_o})}{dt}}{k_f\frac{I}{I_0}} \quad (3)$$

From equation (1) and (2), the $k_b$ and $k_f$ parameter for IAA nanosensor were estimated using *in vitro* kinetic data:

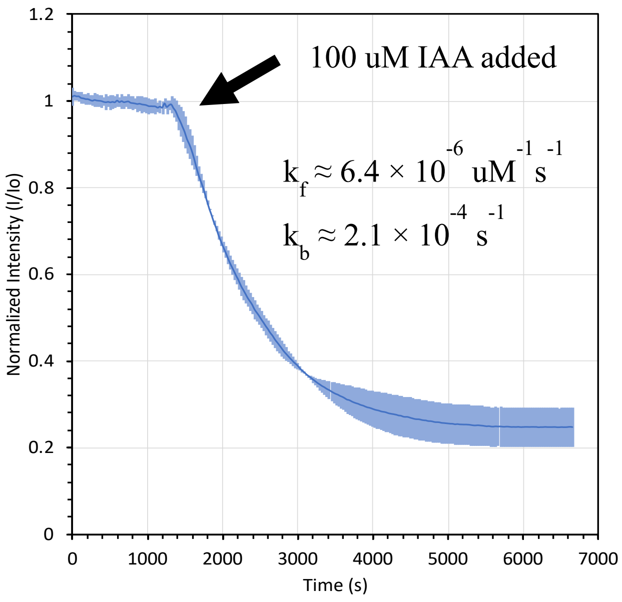

(The solid line represents mean values and the shaded region represents the s.d. (*n*=3).

**One-dimensional steady-state reaction-diffusion model**

For the data in Figure 4c, we develop a model for the spatial IAA distribution from the midrib ($z = 0$) to the margin of the leaf ($z = L$) in the treated side of the leaf. IAA ($C$) is produced with rate $\gamma$ across the induced site with a distribution modeled by a Gaussian function centered about $a$ with standard deviation $\sigma$. IAA is inactivated and reacted away with rate $k_{sink}$ throughout the leaf and is free to diffuse within the leaf with diffusivity $D_f$. The mass balance is as follows:

$$0 = D_f \frac{d^2C}{dz^2} + \gamma \frac{1}{\sigma\sqrt{2\pi}} Exp\left[\frac{-(z-a)^2}{\sigma^2}\right] - k_{sink}C, \quad \text{(S1)}$$

with boundary conditions

$$D_f C'(0) = 0, \quad \text{(S2)}$$

$$D_f C'(L) = -k_c(C(L) - C_\infty) \cong -k_c C(L). \quad \text{(S3)}$$

Here, $k_c$ is a mass transfer rate between the blade of the leaf and the margin of the leaf, and a convective boundary condition is used to model the margin (Equation S3). We assume that IAA concentration is negligible in the margins $C_\infty \cong 0$. Equation S2 represents the midrib of the leaf as an impermeable boundary.

To facilitate analysis, we introduce the following non-dimensional variables:

$$\bar{C} = \frac{C}{C_0}, \qquad x = \frac{z}{L}, \qquad \bar{a} = \frac{a}{L}, \qquad \bar{\sigma} = \frac{\sigma}{L}, \quad \text{(S4)}$$

which yields dimensionless groups

$$\bar{\gamma} = \frac{L\gamma}{D_f C_0}, \qquad \phi^2 = \frac{k_{sink}L^2}{D_f}, \qquad k_{margin} = \frac{k_c L}{D_f}. \quad \text{(S5)}$$

Here, $C_0$ is the initial auxin concentration before induction. Rewriting the dimensionless mass balance yields

$$\frac{d^2\bar{C}}{dx^2} + \bar{\gamma}\frac{1}{\bar{\sigma}\sqrt{2\pi}} Exp\left[\frac{-(x-\bar{a})^2}{2\bar{\sigma}^2}\right] - \phi^2\bar{C} = 0, \quad \text{(S6)}$$

with boundary conditions

$$\bar{C}'(0) = 0, \quad \text{(S7)}$$

$$\bar{C}'(1) = -k_{margin}\bar{C}(1). \quad \text{(S8)}$$

For data analysis, a representative strip of data approximately centered about the induced spot was averaged into one dimension. A Gaussian-weighted moving average filter with a window of 30 was applied to the data.

The Damköhler number can be rewritten as $\phi^2 = \left(\frac{L}{\lambda}\right)^2$, where $\lambda = \sqrt{\frac{D_f}{k_{sink}}}$ represents the length of the reaction zone for IAA. Taking the analyzed region to have $L = 0.48$ cm, and averaging the fitted Damköhler numbers to get $\phi = 3.4$, we calculate $\lambda = 0.14$ cm.
